# A general deep learning model for bird detection in high resolution airborne imagery

**DOI:** 10.1101/2021.08.05.455311

**Authors:** Ben G. Weinstein, Lindsey Garner, Vienna R. Saccomanno, Ashley Steinkraus, Andrew Ortega, Kristen Brush, Glenda Yenni, Ann E. McKellar, Rowan Converse, Christopher D. Lippitt, Alex Wegmann, Nick D. Holmes, Alice J. Edney, Tom Hart, Mark J. Jessopp, Rohan H Clarke, Dominik Marchowski, Henry Senyondo, Ryan Dotson, Ethan P. White, Peter Frederick, S.K. Morgan Ernest

**Author notes:** Open Research Statement The data, code and imagery used in this work are all available as open-source resources with permissive licenses. The code for recreating the analysis is noted in the manuscript (https://github.com/weecology/BirdDetector) and is available through a previously published python package (https://deepforest.readthedocs.io/). The data and models have been uploaded to Zenodo (https://doi.org/10.5281/zenodo.5033174).

## Abstract

Advances in artificial intelligence for computer vision hold great promise for increasing the scales at which ecological systems can be studied. The distribution and behavior of individuals is central to ecology, and computer vision using deep neural networks can learn to detect individual objects in imagery. However, developing supervised models for ecological monitoring is challenging because it needs large amounts of human-labeled training data, requires advanced technical expertise and computational infrastructure, and is prone to overfitting. This limits application across space and time. One solution is developing generalized models that can be applied across species and ecosystems. Using over 250,000 annotations from 13 projects from around the world, we develop a general bird detection model that achieves over 65% recall and 50% precision on novel aerial data without any local training despite differences in species, habitat, and imaging methodology. Fine-tuning this model with only 1000 local annotations increase these values to an average of 84% recall and 69% precision by building on the general features learned from other data sources. Retraining from the general model improves local predictions even when moderately large annotation sets are available and makes model training faster and more stable. Our results demonstrate that general models for detecting broad classes of organisms using airborne imagery are achievable. These models can reduce the effort, expertise, and computational resources necessary for automating the detection of individual organisms across large scales, helping to transform the scale of data collection in ecology and the questions that can be addressed.

## Introduction

Airborne image capture is revolutionizing data collection in ecology by providing information on animal presence, abundance and behavior at unprecedented scales (Weissensteiner et al. 2015, Bondi et al. 2018, Reintsma et al. 2018, Afán et al. 2018). A central challenge in airborne monitoring is converting the large amount of sensor data into ecological information. Ecological image annotation is laborious and the amount of imagery collected can quickly overwhelm human annotators (Kellenberger et al. 2020). Automated tools for animal detection are therefore critical to make image-based data collection feasible at large scales. Computer vision using deep neural networks is a subfield of artificial intelligence that connects sensor data, such as image pixels, to semantic concepts such as the category and location of individual objects within an image. The rapid growth of computer vision for ecological monitoring has allowed surveys to scale to unprecedented extents using ground-based (Berger-Wolf et al. 2017) and airborne imaging systems (Moreland et al. 2015). There have been a number of recent studies demonstrating that with sufficient quantities of human-annotated data, computer vision using deep neural networks can accurately predict animal location, abundance and behavior (Crall et al. 2013, Beijbom et al. 2015, Bowley et al. 2018, Weinstein 2018, Bondi et al. 2018, Kellenberger et al. 2018, Torney et al. 2019, Willi et al. 2019, Beery et al. 2020, Ahumada et al. 2020). However, developing and deploying models for ecological monitoring can be challenging because: 1) large amounts of human labeled training data are typically necessary for training deep neural networks to convergence, 2) building neural networks requires technical expertise and access to significant computational infrastructure that are often unavailable to ecological teams, and 3) the large number of parameters in deep neural networks tend to overfit to available data preventing application across space and time.

One solution to these challenges is the development of generalized models that can be applied to a broad range of species and ecosystems. Generalized models attempt to produce a single model that works effectively regardless of the details of image background and target object (Kawaguchi et al. 2020). Due to shared evolutionary history, animals within broad taxonomic groups, such as class (e.g *Aves, Mammalia*) and order (e.g. *Charadriiformes, Carnivora*) share general visual characteristics that can be useful for detection in natural landscapes. While there will be significant variation in appearance within these groups, the goal is to combine annotations from many studies to produce the large data sets needed to train a general detector. The majority of computer vision models developed in ecology focus on a single species or ecosystem. As a result, it is not clear how accurately deep learning models predict under truly novel ecological conditions, how effective fine-tuning is for producing improved predictions for individual studies, and how these models compare to similar models developed using only training data collected specifically for that study.

The detection of large birds in airborne imagery is an ideal scenario for developing and evaluating the performance of generalized models. Birds are a major indicator of ecosystem health and represent an important intersection of remote sensing and conservation (Gregory and Strien 2010). Airborne monitoring of birds using unoccupied aerial vehicles (UAVs) and airplanes is increasingly common due to the need to monitor birds over large scales (Groom et al. 2013, Dulava et al. 2015, Kim and Kim 2020, Pfeifer et al. 2021). Much of this work involves hand counting birds in imagery using human annotators. When computer vision methods are applied, they typically use between 10,000 and 40,000 local annotations developed for a single species and ecosystem (Chabot et al. 2018, Liu et al. 2020). The widespread use of airborne imagery for monitoring birds has created sufficient data for developing a general model, but doing so is challenging because bird species vary dramatically in appearance and the airborne imagery is not collected in a standardized manner, with each monitoring effort collecting data using different airborne vehicles, sampling protocols, and sensors.

Here we develop and evaluate a deep learning object detection model that identifies large birds in high resolution airborne imagery across species and ecosystems. We build from a well-annotated image monitoring program of long-legged wading birds in the Everglades ecosystem of South Florida and add data from projects around the world to train the model to detect birds regardless of species and habitat (Table 1, Figure 1). To address variation resulting from differences in sensors and survey methods, we used data from a range of acquisition platforms and performed on-the-fly data augmentation changing the sizes of individual annotations to represent variation in the height and resolution of image capture. We show that this model performs well for detecting novel species in novel environments, that its performance can be improved by fine-tuning with only small amounts of human-labeled data, and that these fine-tuned models outperform models based only on local training data. This approach creates a framework for general models covering common animal taxa and will lower the barrier to automated monitoring using airborne imagery.

**Table 1.**
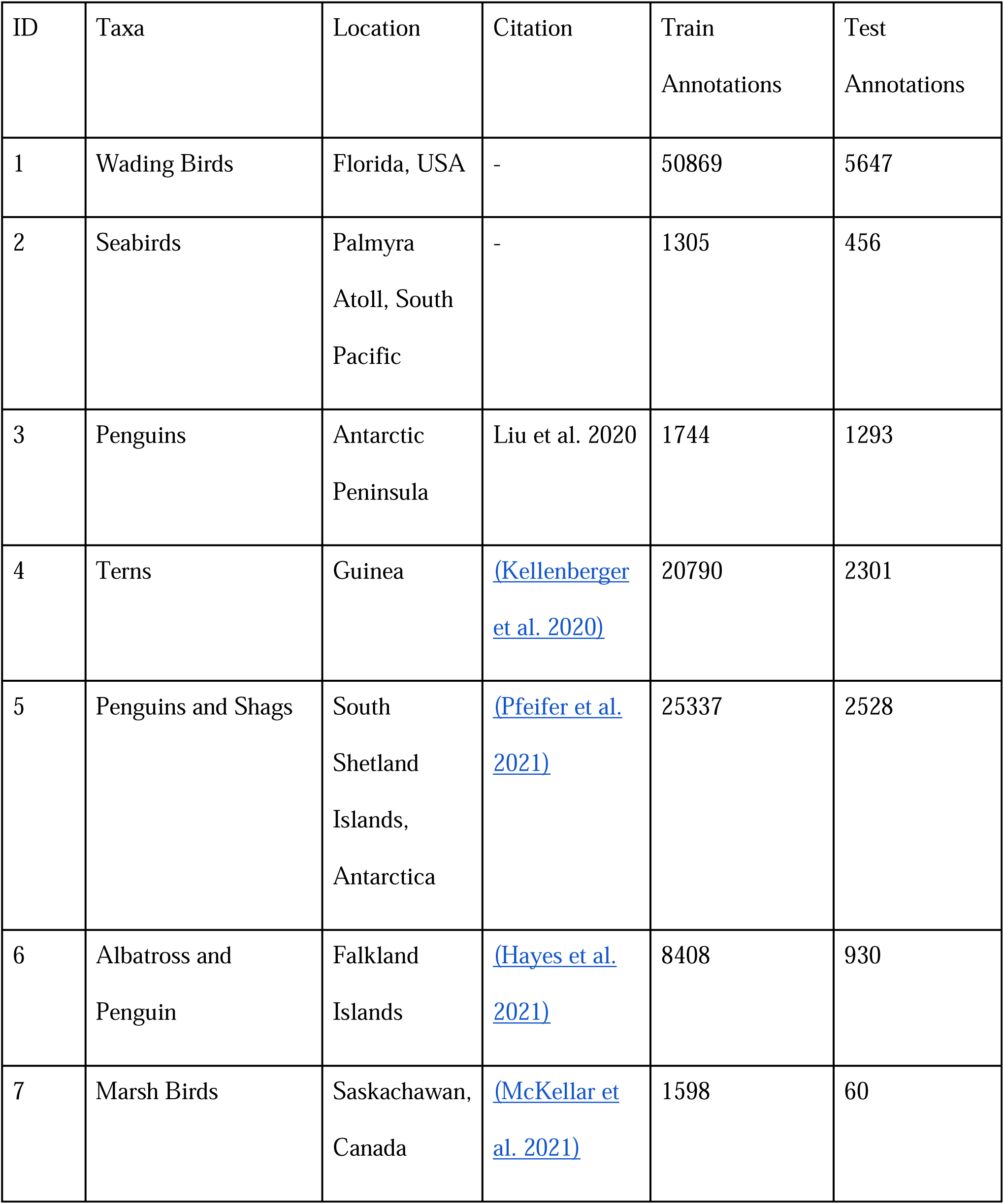
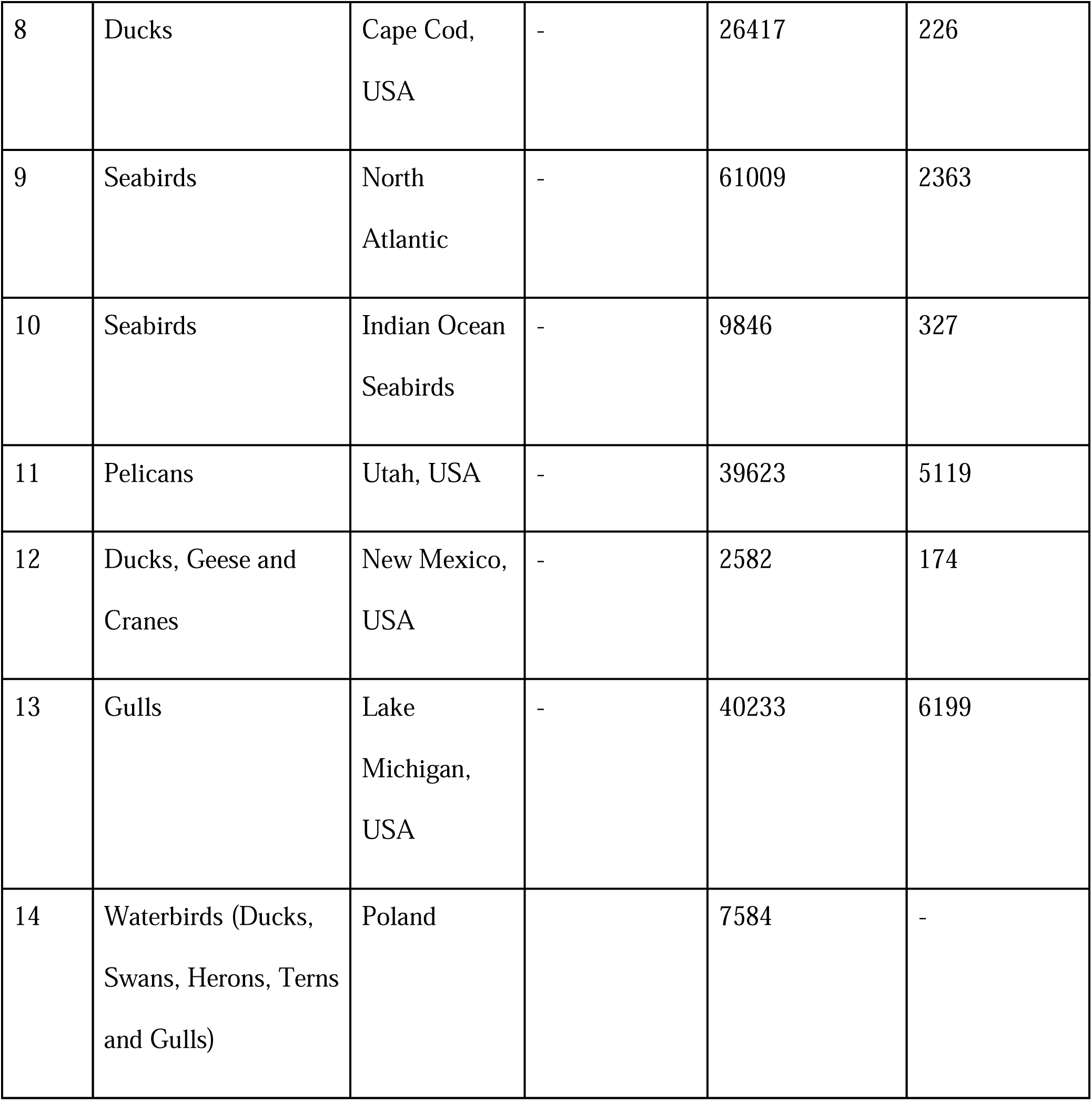
Description of the datasets used to train and evaluate the general model. For more detail on taxa, resolution and acquisition conditions, please see Appendix S1.

**Figure 1:**
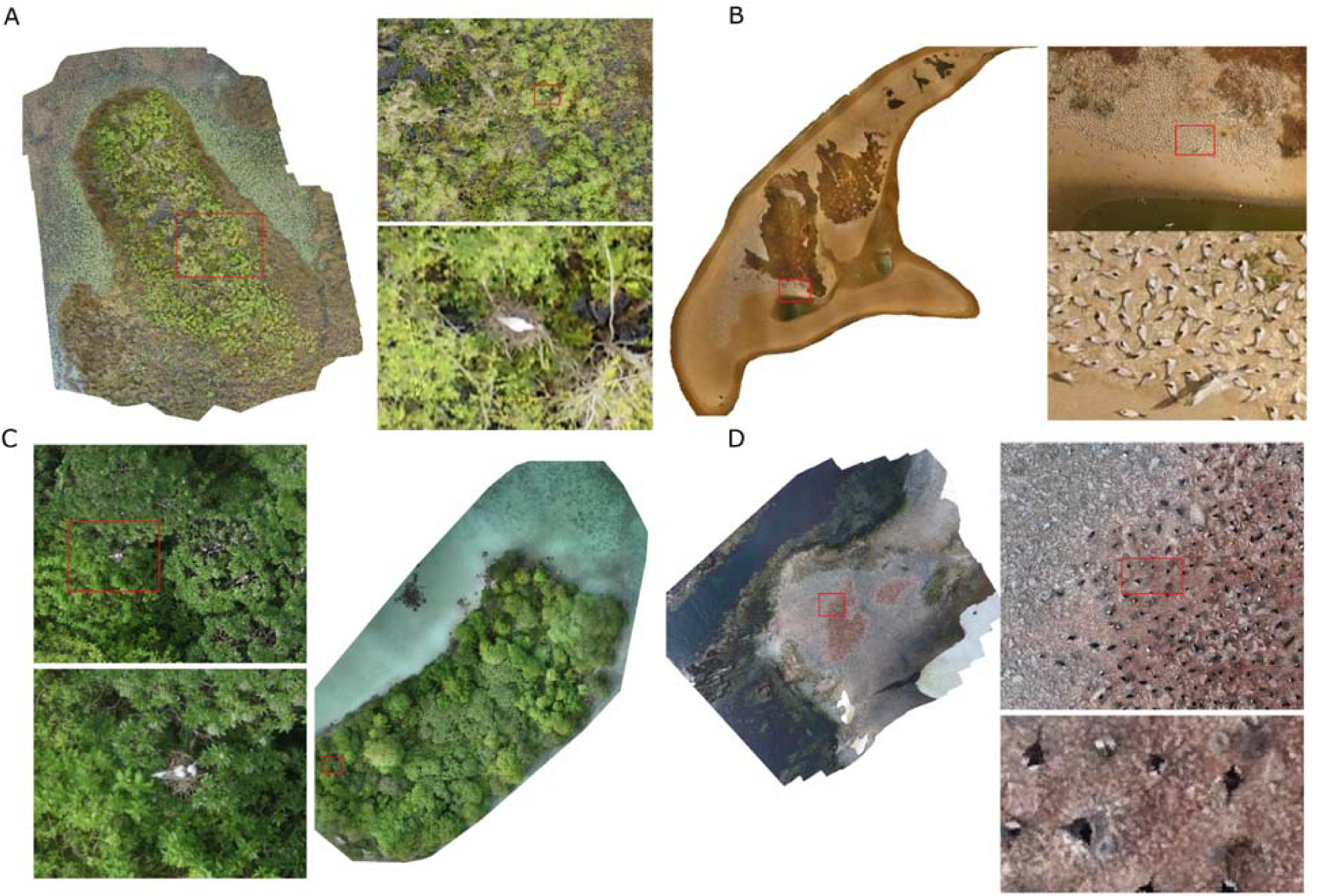
Example orthomosaics for A) the Everglades dataset (ID = 1 in Table 1), B) West African Terns (ID = 4), C) Seabirds from the South Pacific (ID = 2), and D) Chinstrap Penguins from Antarctica (ID = 3). The datasets differ in acquisition backgrounds, target species, individual density and camera specifications.

## Materials and Methods

### Data

Initial model development was conducted with a large dataset of wading bird imagery collected in the Everglades. This data was collected using UAV surveys over wading bird nesting colonies from January -June 2020 to capture roosting and nesting behavior of wading birds in the central Everglades, Florida, United States. We used a DJI Inspire II quadcopter fitted with a Zenmuse X7 RGB camera and 35mm equivalent fixed lens. Flight transects were created using the DJI Ground Station Pro App® and included 80% front and 75% side overlap of imagery collected at 15° off-nadir for a ground resolution of approximately 1cm. Flights were conducted between 76 and 91 m above ground level (AGL) (dependent on disturbance and air space). Target species of wading birds included White Ibis (*Eudocimus albus*), Roseate Spoonbill (*Platalea ajaja*), Great Egret (*Ardea alba*), Snowy Egret (*Egretta thula*) and Wood Stork (*Mycteria americana*), but images also included incidental detections of other species in the ecosystem. After data acquisition and preprocessing, we divided the orthomosaic into small clips and annotated all visible birds with a single x,y point. Points were then transformed into boxes using a fixed buffer size of 0.25m. We annotated a total of 4,653 images yielding 57,000 individual bird annotations. Prior to model training, orthomosaic image tiles were divided into 1500 × 1500 pixel crops and the resulting crops were split into train and test samples. The training data sample for each dataset was composed of randomly selected crops that represented as close to 90% of the annotations as possible within that dataset given the structure of the annotations within the images (range 50% - 90%).

To build a global dataset, we contacted ecologists and conservation biologists using airborne monitoring to request datasets from airborne images with largely nadir camera position and ground sampling distance of 1-3cm (Figure 1). We obtained 13 datasets covering over 250,000 annotations for model training and testing (Table 1). All datasets came from airborne bird monitoring projects with the exception of Seabird Watch (Table 1, ID = 9), which uses ground based remote time lapse cameras to view distant rocky cliffs. This dataset was judged sufficiently similar to be included based on size and resolution. For each dataset we split into training and test sets. Wherever possible test sets were selected to be separate in time or space from training images. For example, if there were multiple flights in an area, each flight would occur only in train or test sets. Or if multiple islands were surveyed, each island would occur only in train or test sets. See Appendix S1 for information on each dataset, including camera specifications, species lists and flight information. Code, processed images and annotations are made available on Zenodo (Weinstein 2021). The final model (https://github.com/weecology/BirdDetector) and evaluation procedures are made available through the DeepForest python package allowing users to extend, train and evaluate models with minimal difficulty (Weinstein et al. 2020).

### Models and Analysis

A general model for bird detection should predict birds in novel environments, allow customization to new datasets using local annotations, and perform better than solely local data. To test these characteristics, we trained a suite of models for analysis (Table 2). All models shared the same architecture and general workflow but differed in input data. The architecture was initially developed for identifying trees in airborne imagery by Weinstein et al., (2020b, 2019). The model was a one-shot object retinanet detector with a convolutional neural network backbone (Lin et al. 2017) implemented in the ‘DeepForest’ Python package (Weinstein et al. 2020a). The retinanet detector uses focal loss to increase the weight of difficult to predict images, reducing the overfitting to easy-to-predict samples. The retinanet backbone was a resnet-50 network pretrained on the ImageNet classification benchmark.

**Table 2.** Model descriptions and terminology for the analysis of the general bird model. Global annotations come from all datasets except for the dataset being evaluated. Local annotations come from the dataset being evaluated. During zoom data augmentation, images are randomly preprocessed to zoom in or out on a subset of birds within an image to mimic flights at different heights. All models were trained using SGD optimization with learning rate of 0.001 and momentum of 0.9 with a batch size of 32 on two NVIDIA DGX A100 GPUs

| Model | Global<br>Annotations | Local<br>Annotations | Zoom<br>Augmentation | Epochs | Description |
| --- | --- | --- | --- | --- | --- |
| Cross-validation | Yes | No | Yes | 12 | For each dataset we trained on all other datasets and then predicted a portion of the withheld dataset. |
| Local-only | No | Yes | No | 70-110 | Individual models using only local annotations for each dataset. This model was repeated for up to 1000, 5000, 10000 and 20000 local annotations. |
| Fine-tune | Yes | Yes | No | 20 | For each dataset we started from the cross-validation model using all other datasets and then fine-tuned using local annotations. This model was repeated for up to 1000, 5000, 10000 and 20000 local annotations. |
| General<br>model | Yes | Yes | Yes | 12 | Final model trained with all<br>training and test data for future<br>use |

One of the major challenges with building generalized models for airborne bird detection is that airborne data from different sources varies in the height of image capture and resolution of the camera. This leads to differences in size, contrast and detail of birds among datasets. To synthesize bird detections at different resolutions we performed on-the-fly-data augmentation during training to change the sizes of individual annotations to represent variation in the height and resolution of image capture (Zoph et al. 2019). During each batch, a random annotation was selected, and then a randomly sized box was placed with this focal annotation at its center. To avoid over zooming on the annotation and filling the entire image, we set a minimum size of 0.15 times the original image size. This technique reduces overfitting but cannot fully remedy differences in image resolution since it is possible to downscale images, but difficult to realistically upscale images. In addition to random zooming, each image was randomly flipped over the x or y axis with a probability of 0.5.

To evaluate the ability of the general model to predict birds in novel locations, we performed a leave-one-out cross-validation analysis. For each large dataset, we trained a ‘cross-validation’ model using all other datasets and then predicted the test images of the withheld dataset (Table 2). While the final ‘general’ model available to users is trained with every dataset, this leave-one-out strategy is a conservative proxy for future use because it represents how well a general model works when not trained on data for a new monitoring effort. Each of the cross-validation models were trained with a batch size of 32 for 12 epochs. Training was performed using stochastic gradient descent (SGD) with a momentum of 0.9 and an initial learning rate of 0.001. Learning rates were reduced by 50% when validation loss had not decreased by more than 0.001 over a period of 10 epochs with a minimum learning rate of 0.00001.

To determine whether starting from the general model improved performance for new datasets, we fine-tuned each ‘cross-validation model’ using local data. For example, to test the ability to customize to the Atlantic Seaduck dataset (ID = 8), we started from the ‘cross-validation’ model trained on all other datasets and added in Atlantic Seaduck annotations. We trained multiple versions of a fine-tuned model, each with a subset of local datasets with 1000, 5000, 10000 and 20000 annotations. We repeated the sub-sampling for the fine-tuned model 3 times to evaluate the effect of image sampling. Finally, to determine whether the fine-tuned model benefited from the pretraining on all other datasets, we trained a ‘local-only’ model that used the same annotations as the ‘fine-tuned’ model, but starting from standard ImageNet weights. The same test dataset was used for the fine-tuned and local-only models. The zoom data augmentation strategy was not used in these models since flight height and object size are largely conserved within each dataset. The local-only models were more sensitive to initial conditions, and given each dataset, we attempted to find the optimal number of epochs that led to model convergence and highest performance.

For all analysis, we used precision and recall on held-out images for model evaluation. The most common evaluation metric in object detection is intersection-over-union, defined as the area of intersection between the true and predicted bounding box, divided by the area of union between true and predicted bounding box. Using this metric, we assessed model recall, defined as the proportion of ground truth boxes correctly overlapping with predicted boxes with an intersection-over-union of greater than 0.2, and model precision, defined as the proportion of predicted boxes which overlap with a ground truth box with an intersection-over-union of 0.2.

We selected this threshold because the vast majority of annotations were automatically created from original points placed on individual birds. The exact outline of individuals is therefore approximate and secondary to the goal of detection and enumeration. To rank models, we also calculate the F1-score for each model, which is a combined score of precision and recall and is calculated as F1 = 2 * (precision * recall) / (precision + recall).

## Results and Discussion

General models for ecological object detection will be most useful if they can detect individuals in novel environments, allow customization to new datasets using local annotations, and produce better detections than models developed with limited local annotations alone. To test for these characteristics, we trained a suite of local and general models for analysis (Table 2). All models shared the same workflow and architecture, a one-shot object retinanet detector with a convolutional neural network backbone (Lin et al. 2017), but differed in input data and evaluation data. We performed on-the-fly-data augmentation during training, zooming in on individual annotations to represent variation in the height and resolution of image capture (Zoph et al. 2019).

To evaluate the ability of a general model to predict birds in novel locations, we performed a leave-one-out cross-validation analysis. For each large dataset, we trained a ‘cross-validation’ model using all other datasets and then predicted the test images of the withheld dataset (Table 2). Each model was judged to have converged by visually assessing the validation loss during training (Figure S3). The mean recall of the held-out dataset was 67.9% (range = 29.8, 95.4%), the mean precision was 52.9% (range = 18.8, 79.7%; Figure 2), and the mean F1 score was 54.5% (range = 30.4%, 81.5%; Table 3). In general, performance was better for datasets with high resolution imagery, such as the West African Terns (ID = 4; recall = 87.7%; resolution = <1cm), whereas lower resolution datasets like the Antarctic Chinstrap Penguins (ID = 5; resolution = > 2cm)) had lower values (recall = 29.8%). Datasets with forested backgrounds similar to the everglades dataset (ID = 1), which forms the backbone of the training annotations, had higher precision, such as the South Pacific Seabirds (ID = 2, precision = 74.0%) whereas datasets with complex aquatic backgrounds had lower precision (e.g Canadian Marshbirds, ID=7, precision = 18.9%). These results suggest that there is the potential for a generalized model to make accurate predictions for completely novel species and environments, but that its performance will depend on having sufficiently diverse data to obtain highly accurate predictions across all novel environments.

**Table 3.**
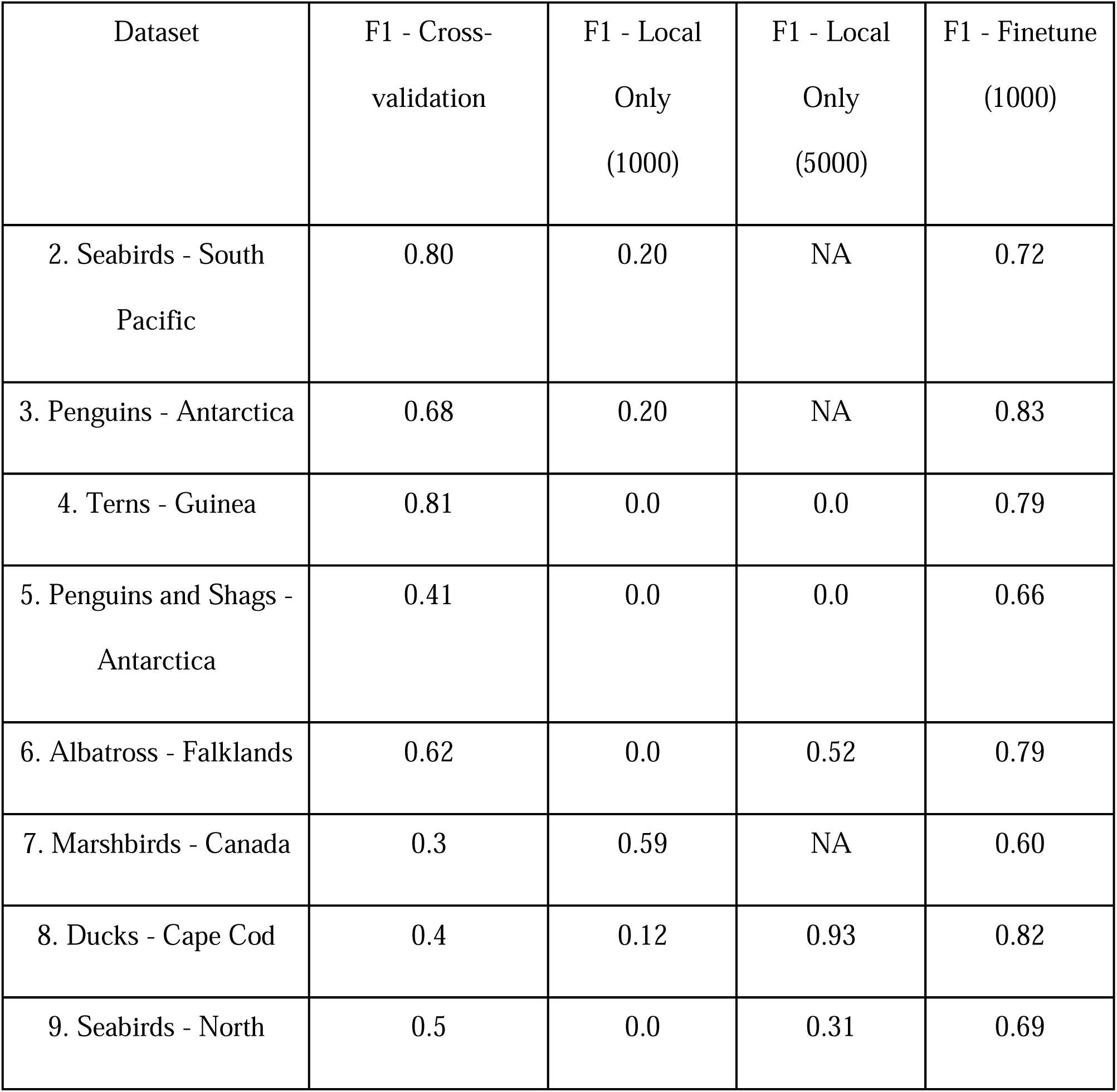
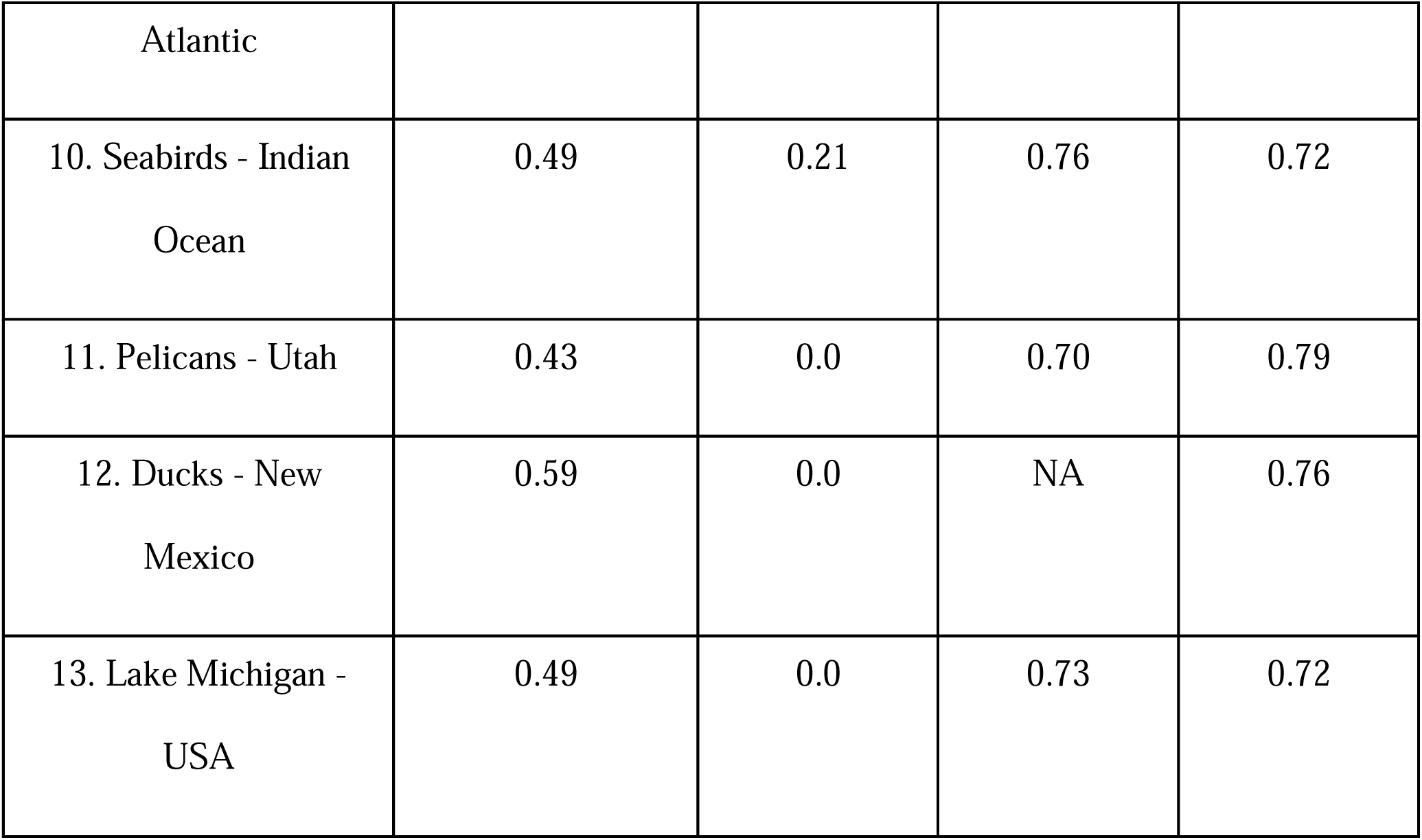
F1 scores for novel datasets. F1 is calculated as F1 = 2 * (precision * recall) / (precision + recall). The cross-validation model is trained on all other datasets and is used to predict the test dataset. The local-only model uses only annotations from the test dataset. The finetune model starts from the cross-validation model and trains on annotations from the test dataset. Two local-only models are shown, one with 1000 annotations and one with 5,000 annotations. Datasets that do not have 5,000 total annotations have NA values.

**Figure 2.**
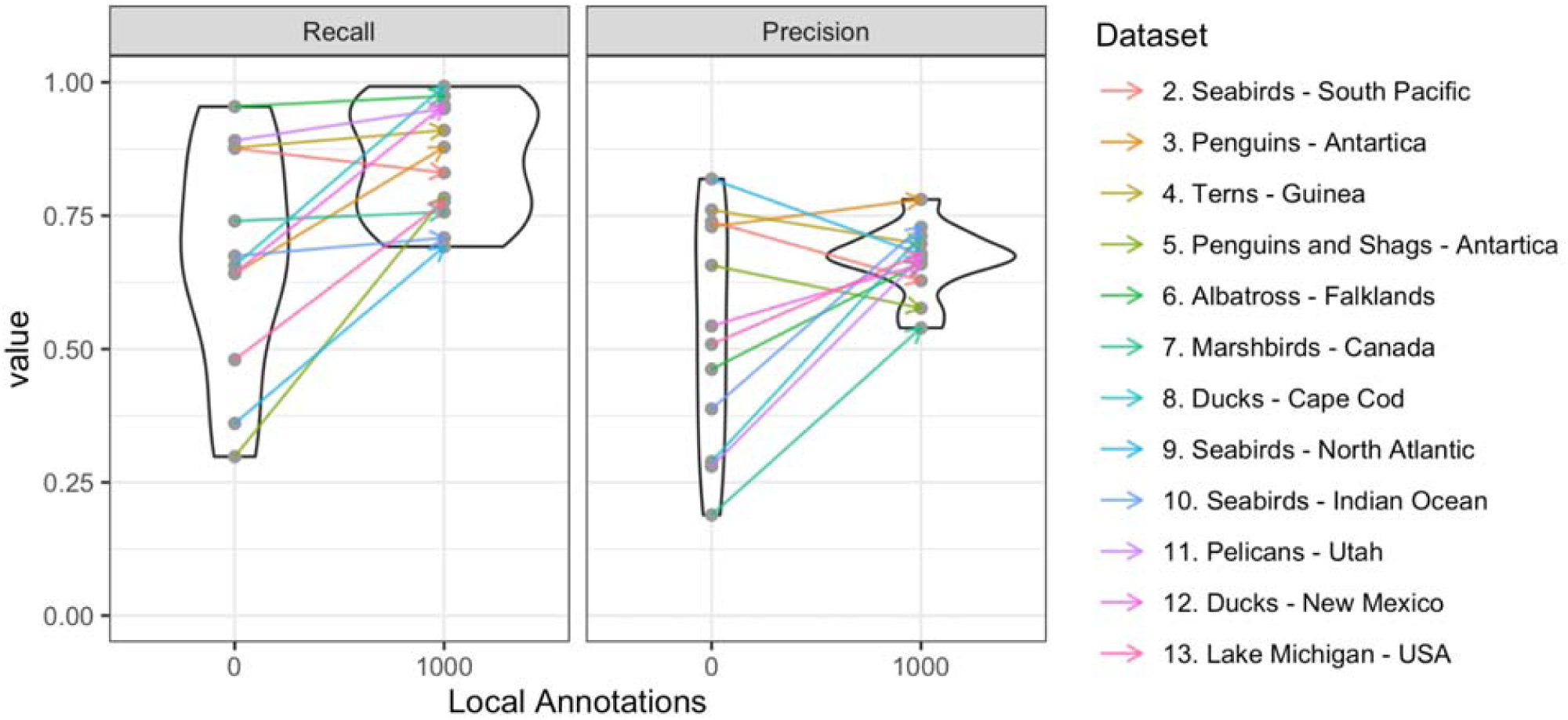
Results of predictions on novel datasets with and without fine-tuning. Recall and precision for each dataset for the cross-validation model with 0 local annotations and the fine-tuned model with 1000 local bird annotations. Recall is defined as the proportion of ground truth boxes that are correctly predicted. Precision is defined as the proportion of predicted that match a ground truth box. The violin plots show the density of points along the Y-axis across all datasets.

General models can be refined to local conditions to improve performance by fine-tuning the model using small amounts of local human-labeled data. Using ∼1000 annotated birds from the local site improved the mean recall, precision, and F1-score to 84.3%, 66.0%, and 74.5% respectively, driven by large improvements in the datasets with the lowest precision and recall scores in the cross-validation models (Figure 2, Table 3). For example, the Atlantic Seaducks (ID = 8) improved from 27% recall to 98% recall (Figure 2A, 2C). Visual inspection showed that improvements often resulted from remedying issues with background objects (e.g, shadows, rocks, leaves) being erroneously predicted as birds, as well as providing the model with information on bird size, which varied due to differences in species, acquisition height, and resolution (Figure 3). Recall and precision where high for all datasets, including those with no clear analog in the general model training data like the SeabirdWatch dataset (where the imagery is from land-based cameras not UAVs; Figure 2C; ID = 9) and the Atlantic Seaducks (where the background is exclusively water; ID = 8). This suggests that general models can be used effectively for most airborne monitoring of large birds using a small number of bird annotations, allowing monitoring efforts to automate bird detection without developing their own deep learning models or investing in time consuming and expensive annotation efforts for every species and environment being studied. For an example of cross-validation prediction for each dataset, see Appendix S1.

**Figure 3.**
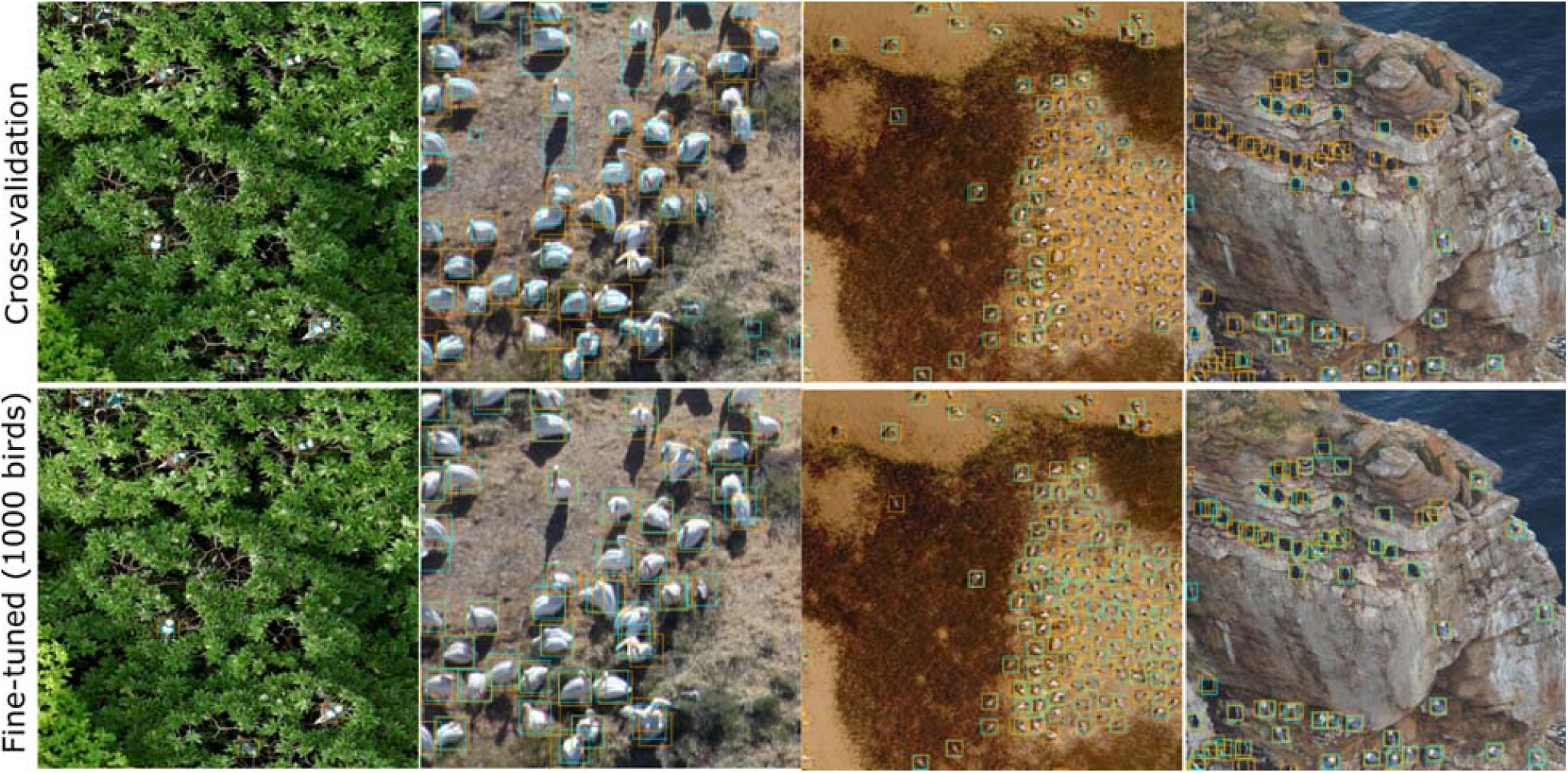
Example images from cross-validation (top) models with no local training data and fine-tuned models (bottom) with 1000 local bird annotations. From left to right, images are from a) South Pacific Seabirds (ID = 2), b) Pelicans from Utah, USA (ID = 11), c) Terns from West Africa (ID = 4), and d) Seabirds from the North Atlantic (ID = 9). Predicted birds are in blue, ground truth birds are in orange.

In addition to making effective predictions with little or no local training data, building on general models may result in more straightforward model development and better predictions for ecological studies even in cases with moderate amounts of training data (Figure 4). This is due to their ability to learn robust general features, thus avoiding overfitting and producing more accurate predictions on images that deviate from the training set, which is a common occurrence when scaling up monitoring efforts. We evaluated the performance of ‘local-only’ models versus the cross-validation models that had the same structure but were initially trained with the training data from all other datasets. In 11 of the 13 test datasets, starting from global model weights improved overall performance (F1-score), often by large margins (Table 3). While this difference was largest when using small amounts of local data, it persisted for some datasets even when using >10,000 local annotations, and the fine-tuned general model always performed at least as well as the local-only model (Figure 4). Local-only models were also highly variable with large changes in performance among runs, sensitive to learning rate and training hyperparameters, and required more computationally intensive training. Fine-tuned models required only 20 epochs of training, whereas local-only models needed to be trained for at least 70 epochs to produce reasonable results (Table 2). Even among local-only models there was large variation in the amount of training needed. For example, the West African Terns dataset (ID = 4) had 0% recall and 0% precision after 70 epochs even when using 20,000 local annotations. Extending to 110 epochs resulted in a rapid increase to 84% recall and 87% precision, but the potential for good predictions for datasets like this would often be missed given the consistently poor performance at shorter training times. This idiosyncratic behavior was difficult to anticipate since the tern dataset is similar to other datasets in terms of the density of birds, image resolution, and background complexity.

**Figure 4.**
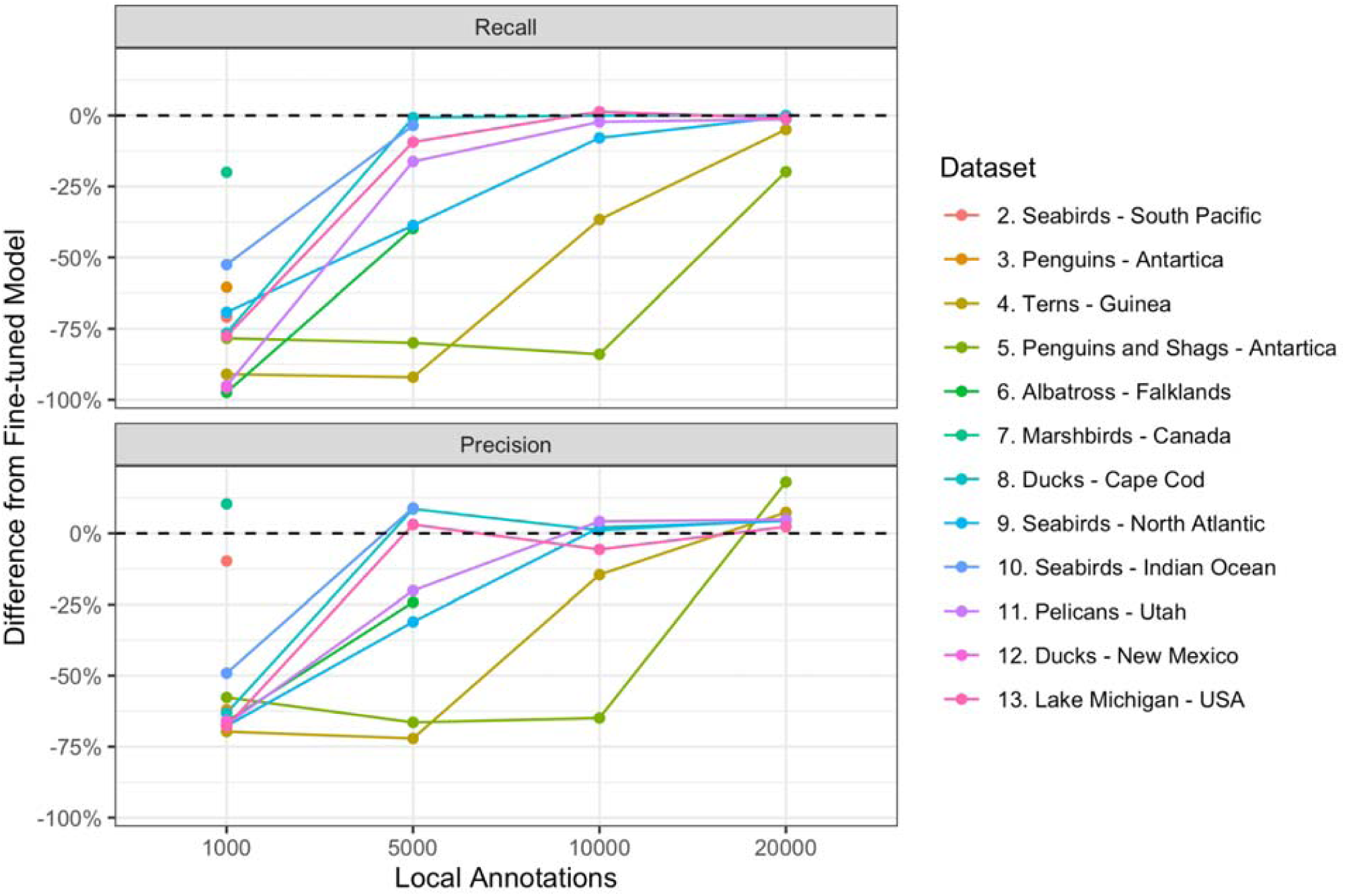
Difference between models fit using only local annotations (the ‘local-only’ model) and general models retrained with the same local data (the ‘fine-tuned’ model) for each dataset using different numbers of local annotations for training. The fine-tuned model is pre-trained on all other datasets, whereas the local-only model contains no annotations from other datasets. Each dataset was run up to its available number of annotations.

Compared to local-only models, the fine-tuned models were more uniform, exhibiting significantly less between-run variance (Figure S2), and achieved good prediction with less data and less intensive computational training. This highlights that even when large numbers of annotations are available for an ecological study, starting with a general model makes developing models easier and provides more stable results. As a result, starting with a general model requires less expertise, less time spent on model development, and fewer computational resources. Our experience developing these models suggests that the difference is significant enough that starting with a general model will allow more studies to successfully build automated approaches to detect individual organisms in airborne imagery and thereby reduce the number of efforts that are limited in the scale of research due to manually annotating all birds being studied.

From a computer vision perspective, one of the major challenges for developing this type of general model for ecological systems is the range of sizes and representations of objects in images caused by differences in the altitude of image acquisition and the sensor resolution. Due to differences in species behavior, UAV regulations, and research questions it is unlikely that there will ever be a standard approach to the acquisition of airborne monitoring imagery in ecology. Therefore, methods are needed that can help overcome this key source of variation. We used a data augmentation strategy to synthesize datasets at different flight elevations and resolutions by zooming in and out on each annotation during model training. This strategy reduced the size-based errors of the dataset, but it did not eliminate them (Appendix S1: Figure S10). Additional strategies to address this challenge going forward include synthesizing new data by combining abundant background data in airborne datasets with bird images from high quality benchmarks such as iNaturalist, which includes tens of thousands of zoomed in images of birds taken by ground-based observers (Van Horn et al. 2018). Synthetic data has been effective in animal detection in thermal airborne imagery and would allow greater control of the range of image scales shown during the model training (Bondi et al. 2018).

While the datasets used in this study differed in capture altitudes, angle, and sensor specifications, they were still broadly similar in using RGB data with resolutions <3 cm. Generalizing to spatial resolutions >3 cm and using non-RGB remote sensing (e.g., hyperspectral imagery) requires further study across sensors and data acquisition strategies. For example, fixed-wing aircraft surveys covering hundreds of miles are unlikely to capture images at ultra-high resolution due to storage and processing limitations. It is unknown how well the features learned in 2 cm imagery will transfer to 10 cm airborne imagery or high-resolution satellite imagery (∼30cm). One approach to this type of generalization is to reduce the higher resolution data and train a series of models to bridge the features learned from high resolution to low resolution data. This is known as ‘curriculum learning’ ((Graves et al. 2017) and can be useful in transferring information among spatial resolutions.

## Conclusion

Aerial imagery is a powerful tool for studying species and ecosystems at temporal frequencies and spatial extents that are difficult using traditional methods, but it comes with computational and analysis challenges that have limited its widespread application. General computer vision models provide a solution for simplifying the processing of aerial imagery to allow researchers to more easily, efficiently, and accurately extract ecological data from large amounts of imagery.

We showed that general models can provide accurate predictions in novel ecosystems and with novel species, with either no local training data or by retraining with very small numbers of annotations. Even when large amounts of local-data are available, starting with general models produces more stable results, with less computational expense, and often performs better than local-only models because of the general features these models learn from other ecosystems and taxa. The ability of general computer vision models to make accurate predictions in novel circumstances will make them an essential tool for monitoring dynamic ecosystems, where species and habitats may change over time or space. Because the need for local hand-annotations is limited, general models can potentially be rapidly deployed in new environments and support aerial monitoring of rare species which can be difficult to study and have limited annotations available for local model development. By reducing the effort, expertise, and computational resources necessary to develop computer vision models for image processing, general models have the potential for revolutionizing the types of data ecology can collect.

Our approach to developing a general computer vision model for bird detection was based on the concept of transfer learning, where a model trained for one task is applied to a different task (Pan and Yang 2010). While we used transfer learning to generalize among bird monitoring projects with new species, in new ecosystems, using images of differing resolutions and image capture features, transfer learning can potentially be applied more broadly to create general detectors across taxonomic groups, including birds, mammals, reptiles, or any other organism visible in airborne images. Using the same general model structure, (Weinstein et al. 2020) developed detection models for individual trees in airborne imagery, suggesting that the underlying architecture is suitable for a broad array of ecological objects. The increased use of UAV-based monitoring, combined with developments in very high resolution satellite imagery (LaRue et al. 2017), increases the potential for animal detection at broad scales with high temporal frequency (Willi et al. 2019). We anticipate that a cross-taxon model of animal detection is achievable with sufficient annotations and research into the most effective way of overcoming the large differences in resolution, object appearance and scale. A general model for automated airborne monitoring of many types of individual organisms would fundamentally change the scales at which we can monitor ecological systems, allowing us to tackle critical questions about the processes that determine population dynamics and biodiversity patterns across scales.

## Acknowledgements

We thank many researchers for contributing published data and images to modeling training including John Neill, Christian Pfeifer, Kyle Landolt, Heather Lynch, Madeline Hayes, Rodrigo Valle, Benjamin Kellenberger, Gary Andrew and Francie Cuthbert. Dan Morris was instrumental in hosting data and organizing the AI4Earth community. Thanks to Mark Koneff and the Branch of Migratory Bird Surveys for their assistance in gathering data and providing insight on model application.

