## Supplementary material for "A general deep learning model for bird detection in high resolution airborne imagery": SI

### Appendix S1

The following is a brief description of the datasets used for training the general model. The ID number corresponds to Table 1. The everglades dataset (ID = 1) is included in the main manuscript.

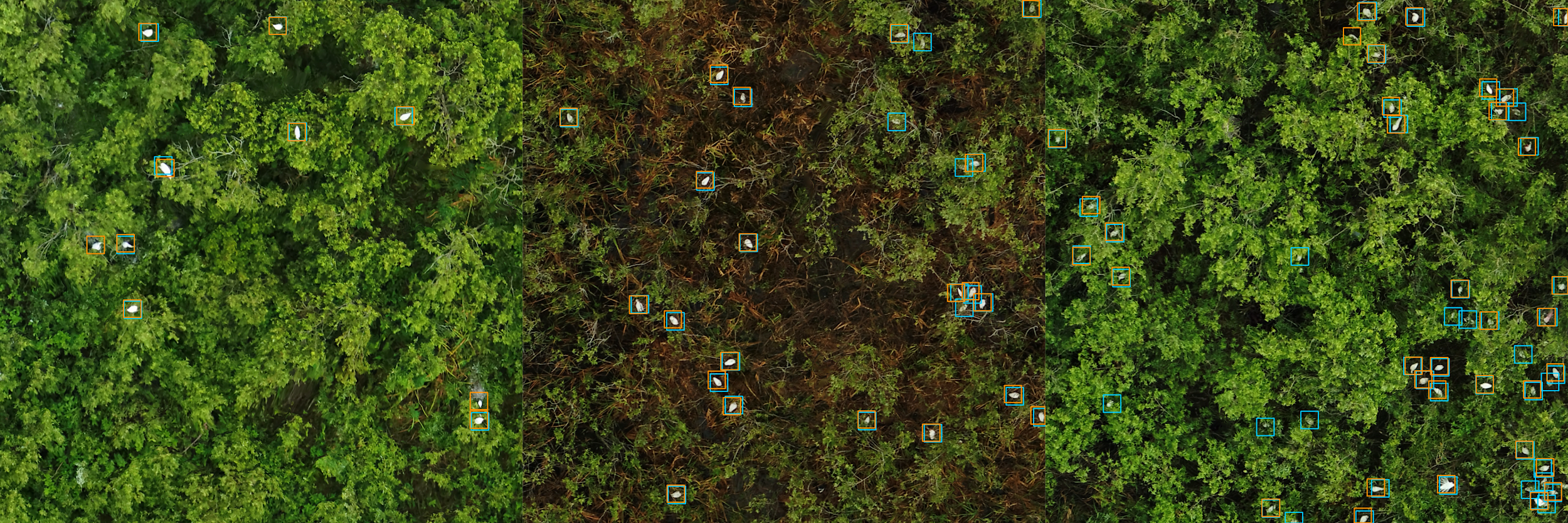

Figure S1. Example predictions from everglades validation images from the preliminary base model using everglades only data. Human-annotated ground truth in blue with machine learning prediction in orange.

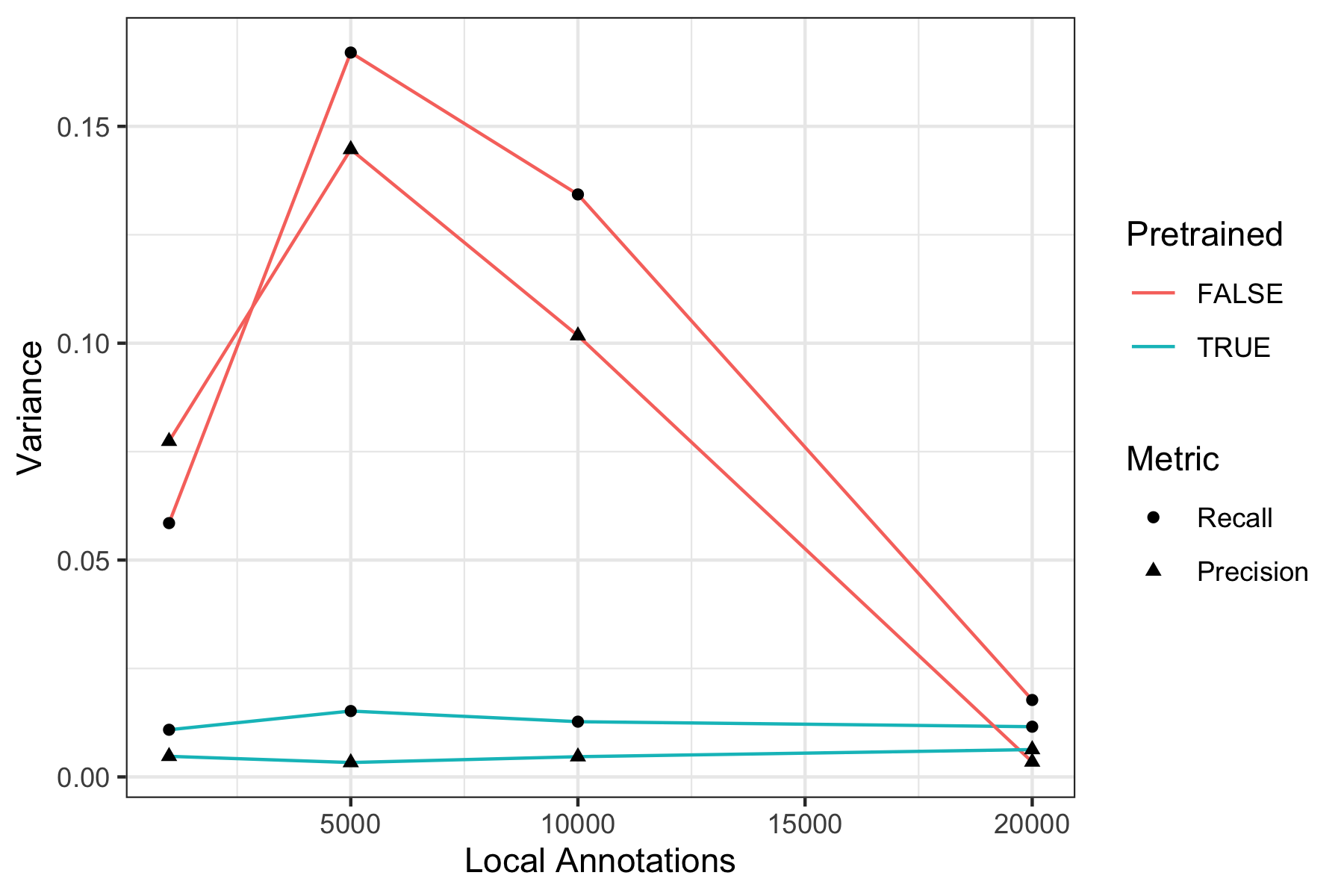

Figure S2. Variance in precision and recall for the fine-tuned (pretrained = True) or local only (pretrained = False) models across all datasets.

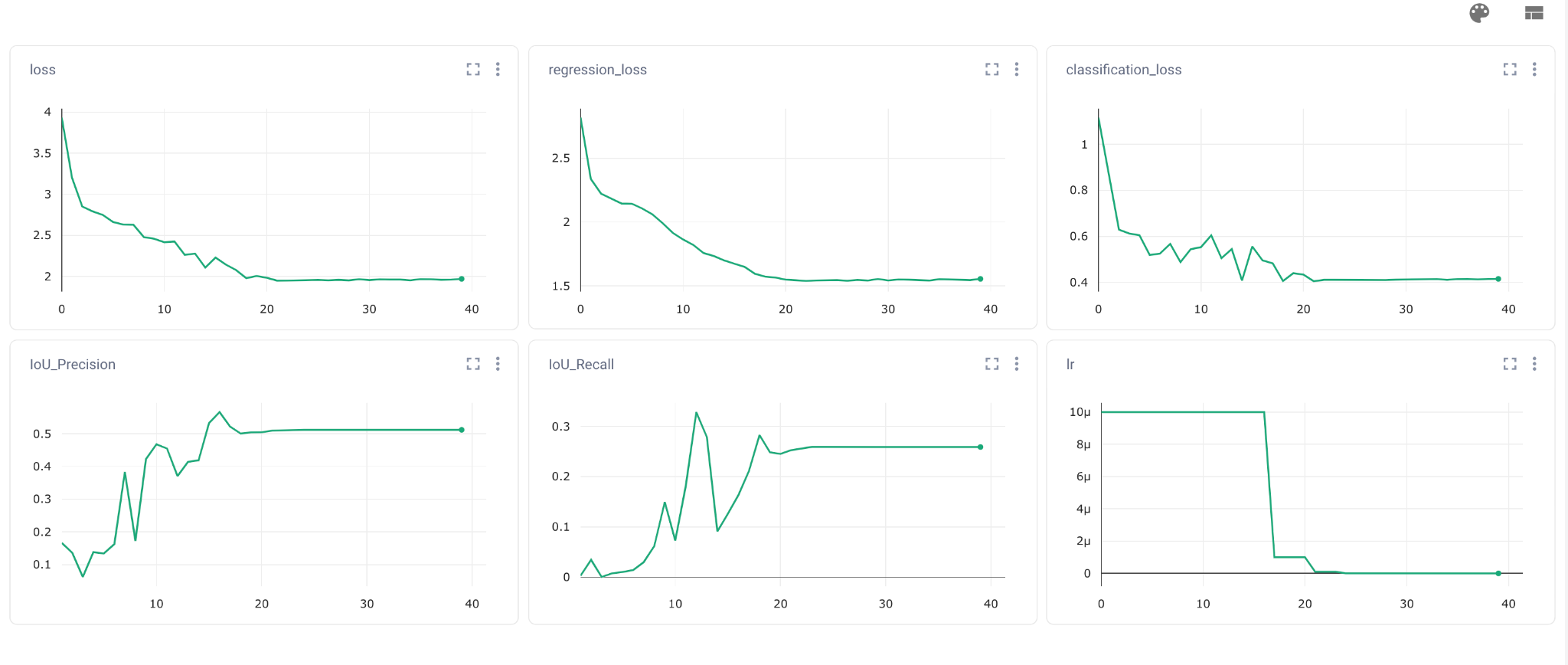

Figure S3. Sample training loss curves for the local-only models for the penguins dataset (ID = 3) run for 40 epochs. The train loss is a composite loss made up of the regression loss, which measures bounding box accuracy, and classification loss, which measures foreground-background detection. The intersection-over-union precision and recall for validation annotations are monitored during training. When validation scores fail to increase during successive epochs, the learning rate (lr) is reduced.

#### 2. Tropical Pacific Seabirds

In 2020, The Nature Conservancy (TNC) collected UAV image data at Palmyra Atoll National Wildlife Refuge in the Northern Line Island Archipelago, Pacific Ocean (5°53′1″N, 162°4′42″W) as a baseline, to measure seabird use of forest canopy habitat following replacement of introduced coconut palms (*Cocos nucifera*) with native tree species. Tropical tree-nesting seabirds show preference for roosting and nesting in broad-leafed tree crowns over coconut palm crowns (Young et al. 2009). By replacing *C. nucifera* with native broad-leaf tree species, TNC and partners aim to increase the spatial distribution of beneficial nutrients transferred to island and nearshore habitats by seabird guano deposition (McCauley et al. 2012). Flights were performed over Palmyra Atoll’s islets in 2020, targeting to observe breeding and roosting behavior of the most abundant tropical seabird on Palmyra, the Red-footed Booby (*Sula sula*) using a WingtraOne fixed-wing VTOL drone with a Sony RX1Rii 42MP full frame with a fixed 35mm lens camera. We used the PPK geopositioning system to improve our location accuracy & eliminate challenges with establishing ground control points in remote, forested survey areas. Flights were performed at 53m with a ground sampling resolution of 0.7cm.

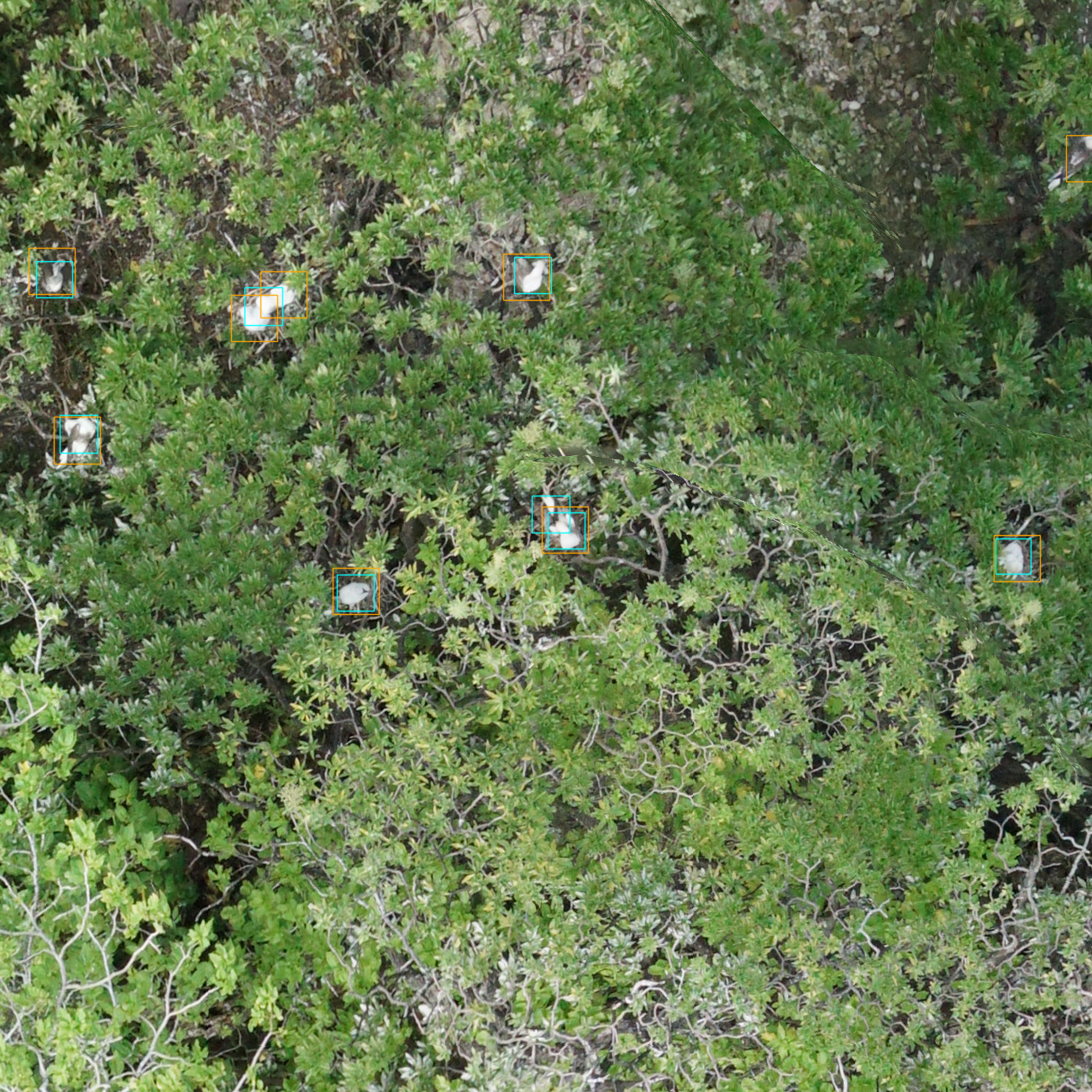

Figure S4. Sample predicted image of birds from the Seabirds from Palmyra Atoll dataset using the cross-validation model. Predictions in blue, ground annotations in orange.

#### 3. Antarctic Penguins from Liu et al. 2020

The Antarctic dataset comes from (Liu et al. 2020), which presented an automated pipeline for identifying post-breeding Chinstrap Penguins (*Pygoscelis antarcticus*) from islands off the Antarctic Peninsula. Surveys were conducted in 2020 using a DJI Phantom 4 quadcopter at altitudes ranging between 25 to 40m with a ground resolution of between 1-2 cm. From the 7,000 labels in Liu et al. 2020 we selected two images to serve as train (‘offshore_rocks_cape_wallace_survey_4’, n=711 penguins) and test (‘cape_wallace_survey_8’, n=744 penguins). Each tile was cut into 900x900 pixel crops to reduce memory constraints. One important distinction for this dataset is that not all penguins that are visible are annotated. Liu et al. (2020) focused on post-breeding individuals and relied on experienced annotators to differentiate breeding status among co-occurring penguins. Model precision can not be assessed because it is not possible to distinguish between predictions which are not penguins, and correct penguins which are not annotated. This distinction also has possible negative effects for model training, since the model will learn to ignore penguin features in the non-annotated individuals. Nevertheless, this difference in annotation protocol is common in ecological datasets and represents a realistic test-case for model transferability.

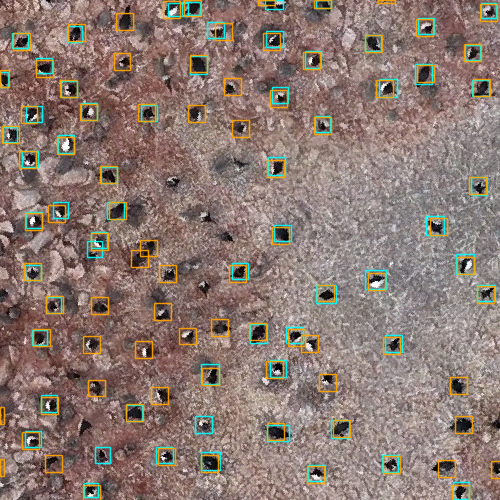

Figure S5. Sample test image from the chinstrap penguins from the Antarctic Peninsula dataset. Predictions from the cross-validation models are in blue, ground annotations are in orange.

#### 4. West African Terns

Kellenberger et al. (2021) conducted UAV surveys of African royal terns (*Thalasseus maximus*) at known breeding sites along the coast of West Africa, between Mauritania and Guinea, in May of 2019. Several other avian species breed at these sites and were included in the count: Caspian terns (*Hydroprogne caspia*), slender-billed gulls (*Chroicocephalus genei*) and grey-headed gulls (*Chroicocephalus cirrocephalus*). The model was unable to distinguish between slender-billed and grey-headed gulls during initial testing, leading the two to be grouped into a single class. Images were acquired by flying a DJI Phantom 4 in parallel transects, at speeds of 3-5 m/s and altitudes ranging between 20 m and 50 m. A 20 Mp camera with a 1” CMOS sensor and 84° field of view was set to take a picture every three seconds. Six RGB orthomosaics were produced, five of which were used for model training with the remaining orthomosaic used for final testing. The test orthomosaic was divided into 274 separate tiles of 800 x 600 pixels laid out in a grid. Point labels were assigned using AIDE and all birds in the image were annotated. The five training orthomosaics were divided into random subset patches and species were hand-annotated through QGIS, with the goal of at least 200 annotations per species.

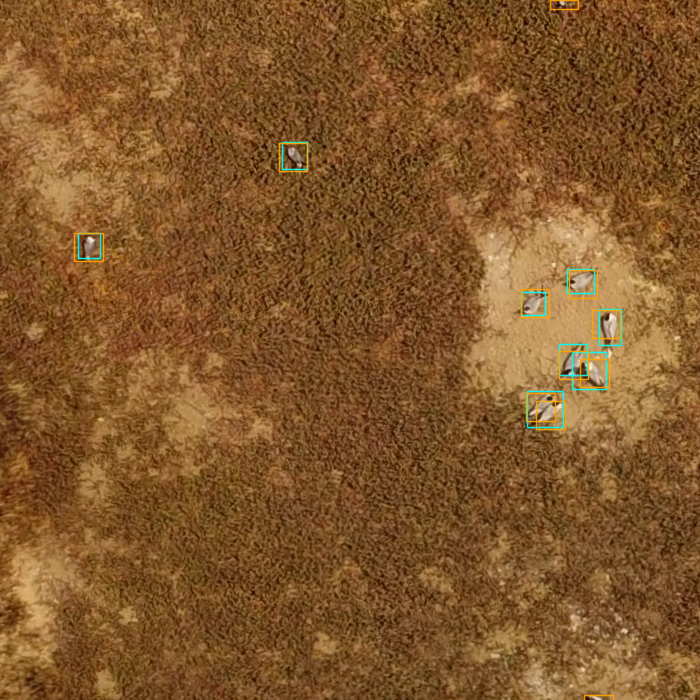

Figure S6. Sample test image from the terns from West Africa dataset. Predictions from the cross-validation model in blue, ground truth annotations in orange.

#### 5. Antarctic Penguins and Antarctic Shags from Pfeifer et al. 2021

Pfiefer et al. 2020 surveyed chinstrap penguin (Pygoscelis antarcticus) and Antarctic Shag (Leucocarbo bransfieldensis) colonies in the South Shetland islands, Antarctica in Dec 2016 using a fixed wing UAV with a light weight (64 g) MAPIR Survey-2 RGB (16 megapixel) digital camera. Images were shot vertically downwards in JPG format every 2.5 seconds in manual exposure mode (f/2.8, shutter speed 1/500 s, ISO 50) at 30–100 m above ground level resulting in a ground sample distance (GSD) of 1–3.4 cm pixel-1. The birds are in dense colonies and often obscured due to rocky backgrounds. The ground sampling distance varies among islands due to topography and flight conditions.

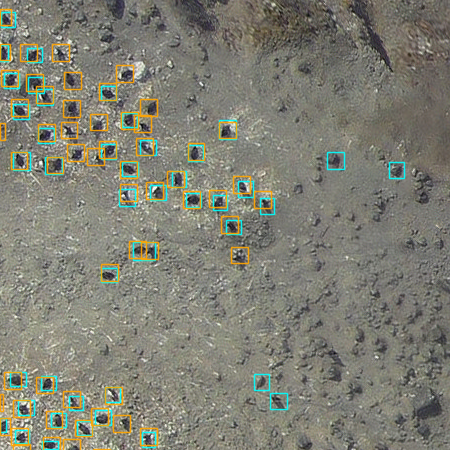

Figure S7. Sample test image of birds from the Penguin and Shag dataset from the South Shetland Islands, Antarctica. Predictions from the cross-validation model are in blue, ground truth annotations in orange.

#### 6. Black-browed Albatross from the Falkland Islands

Hayes et al. (2021) provided a dataset of Black-browed albatross (*Thalassarche melanophris*) and Southern Rockhopper Penguin (*Eudyptes c. chrysocome*) colonies in the Falkland (Malvinas) Islands. Drone surveys of Steeple Jason Island and Grand Jason Island, composed of rocky shores, grassy slopes and steep cliffs, were conducted in November of 2018 and 2019 during the average incubation period for both species. A DJI Phantom with a fixed 9-mm lens and 4,864 x 3,648-pixel resolution was flown at altitudes between 60 m and 90 m depending upon the terrain of the islands as well as to test the lowest resolution threshold. Flights lasted between 20-25 minutes and were run in overlapping parallel lines with patterns defined in DroneDeploy. Pix4D was used to process the images and create orthomosaics with ±1 m horizontal accuracy. Ghosting and edging occurred and were manually edited to present a clean image which was then split into smaller tiles. All birds of both species were manually annotated using VGG Image Annotator, with a 60-pixel overlap for Black-browed Albatrosses and 30-pixel overlap for Southern Rockhopper Penguins.

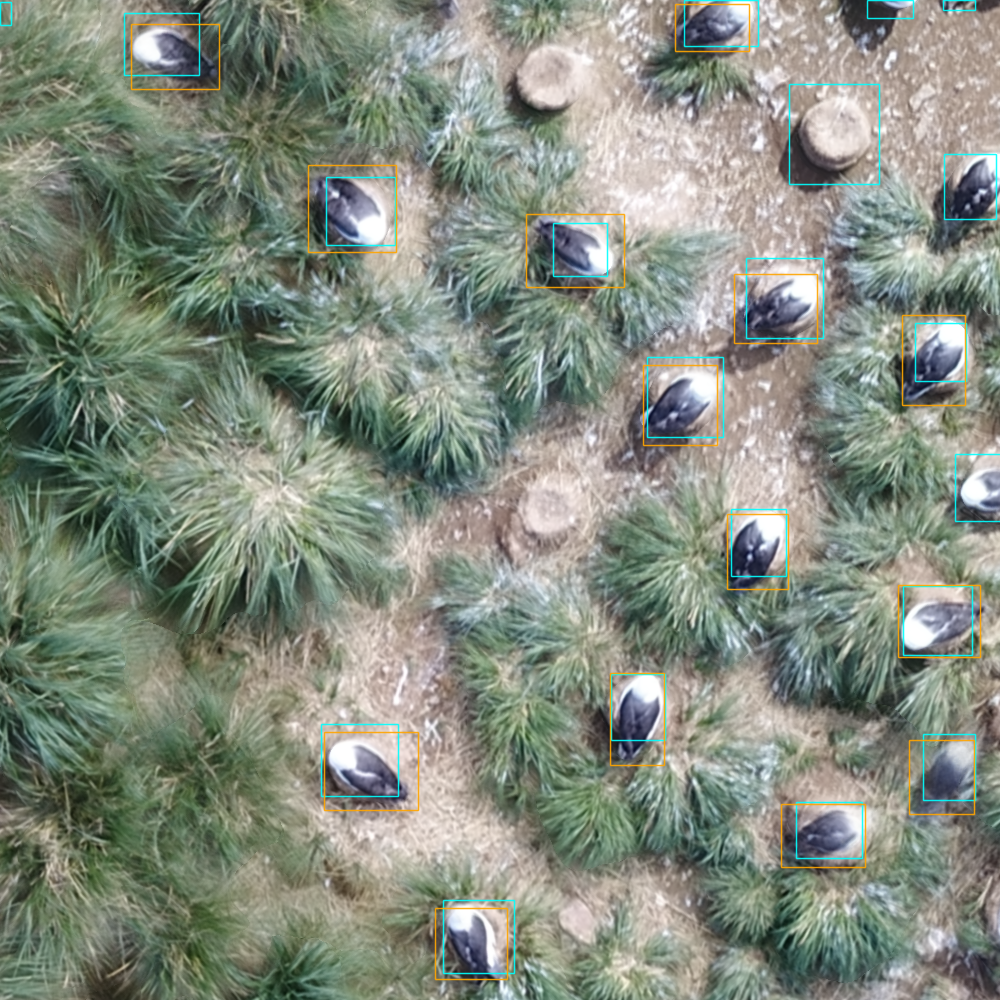

Figure S8. Sample test image of albatross from the Falkland Islands dataset from Hayes et al. 2021. Predictions from the cross-validation model are in blue, ground truth annotations are in orange.

7. Canadian Marsh Birds

In 2019, McKellar et al. (2021) conducted a series of UAV flights to count colonially-nesting marsh bird species in Saskatchewan, Canada. They made use of both thermal (6–16 cm/pixel) and high-resolution visible (0.8–2 cm/pixel) sensors in order to detect and count the nests of target species, and compared UAV-based counts to on-the-ground nest counts. Flights were conducted using a ~9-kg octocopter based on the DJI S1000+ airframe at altitudes ranging from 45 to 120m. The current study made use of orthomosaics from four mixed-species colonies (altitudes of 45-60m). McKellar et al. only annotated nests of five target species. In contrast, for the current study, we re-annotated orthomosaics to include 1) all bird species in addition to target species, and 2) birds on nests as well as birds off nests. Target species included western grebe (*Aechmophorus occidentalis*), Franklin’s gull (*Leucophaeus pipixcan*), black tern (*Chlidonias niger*), Forster’s tern (*Sterna forsteri*), and black-crowned night-heron (*Nycticorax nycticorax*), although colonies including the latter were not included in the current study due to the presence of few birds and mostly empty nests. Other incidental species annotated for the current study include mallard (*Anas platyrhynchos*), ruddy duck (*Oxyura jamaicensis*), unidentified duck spp., American coot (*Fulica americana*), American avocet (*Recurvirostra americana*), yellow-headed blackbird (*Xanthocephalus xanthocephalus*), eared grebe (*Podiceps nigricollis*), horned grebe (*Podiceps auritus*), and red-necked grebe (*Podiceps grisegena*).

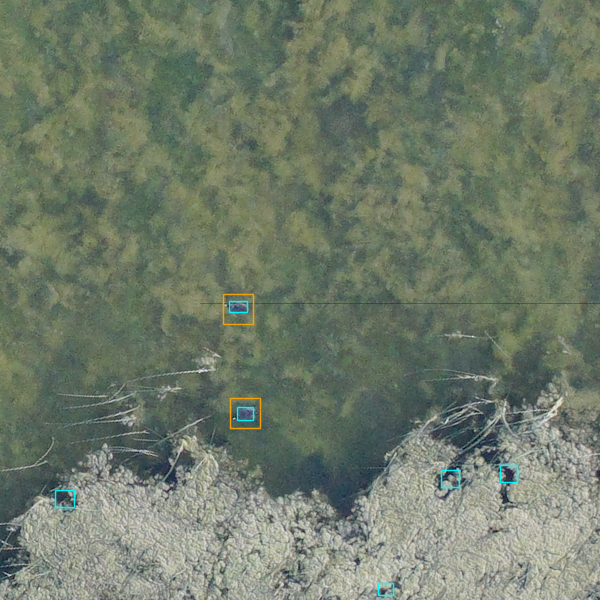

Figure S9. Sample test image from the Canadian Marsh bird dataset from Mckellar et al. (2021). Predictions from the cross-validation model are in blue, ground truth annotations are in orange.

#### 8. Atlantic Seaducks from Cape Cod, USA

Data was collected in February 2017 using aerial imagery of the Nantucket Shoals, Massachusetts, USA acquired by the U.S. Fish and Wildlife Service. The Nantucket Shoals area is a shallow bank with bathymetric, substrate, and tidal characteristics that concentrate prey favored by wintering marine birds and the area can harbor large aggregations annually. All imagery was acquired from a Partenavia P68 fixed-wing airplane using a PhaseOne iXU-R 180 forward motion compensating 80-megapixel digital frame camera with a 70 mm Rodenstock lens. The PhaseOne sensor was integrated with a Global Positioning System and Internal Navigation System in a direct georeferencing system capable of estimating frame-specific exterior orientation parameters necessary for image orthorectification without ground control. The charge-coupled device imager for the PhaseOne camera was 10328 × 7768 pixels. Altitudes of acquisition were between 24.4 to 198.1 m for the Nantucket Shoals site and 21.3 to 42.7 m for the Lake Michigan site. Ground sample distance (i.e., pixel resolution) ranged from 0.18 to 1.47 cm for the Nantucket Shoals site and from 0.14 to 0.32 cm for the Lake Michigan site. Imagery was collected in Phase One’s proprietary IIQ raw image format and converted to color-balanced TIFF images using Phase One’s Capture One image processing software.

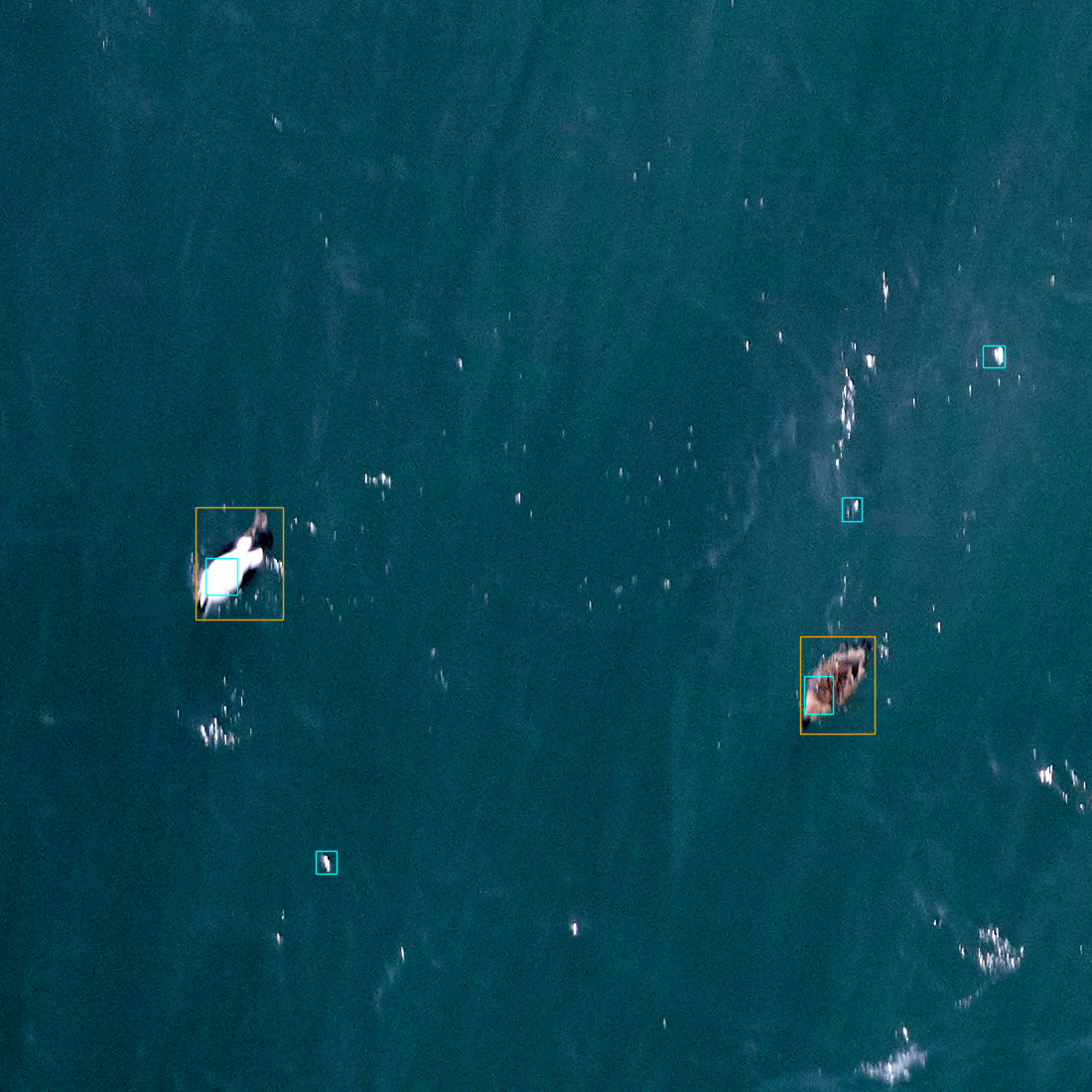

Figure S10. Sample test image from the Atlantic seaduck dataset. Predictions from the cross-validation model are in blue, ground truth annotations are in orange.

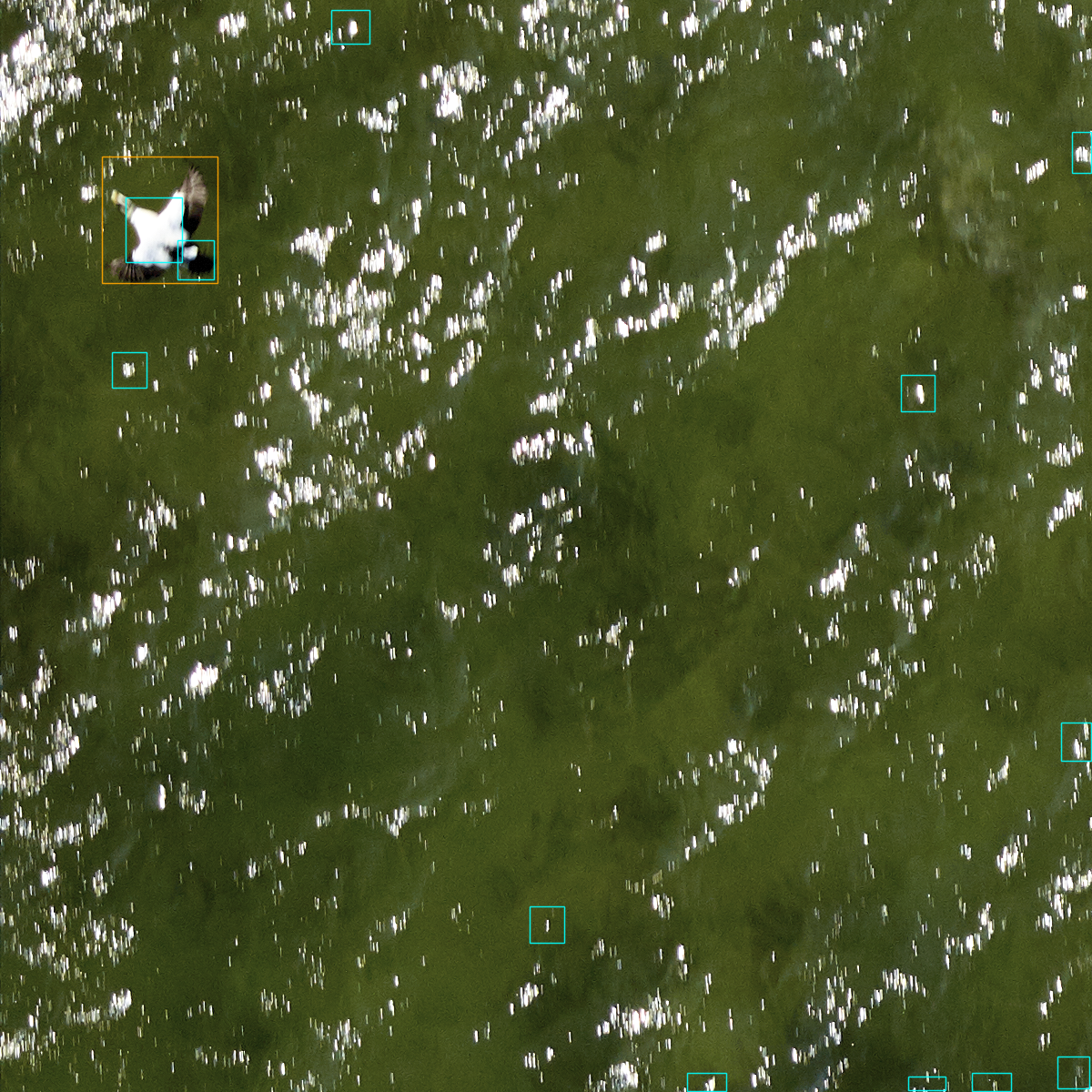

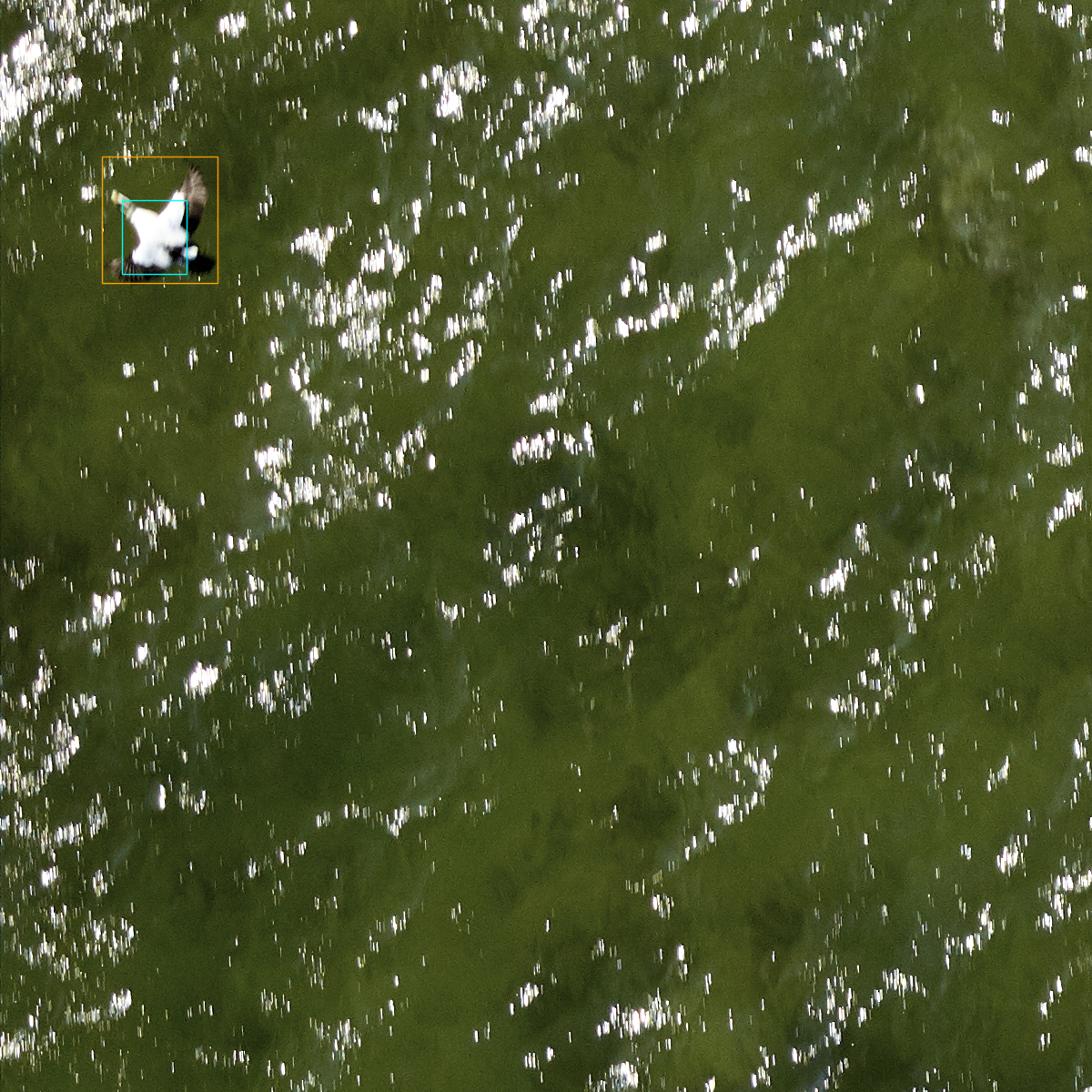

Figure S11. The importance of data augmentation for adjusting to bird detections at a variety of image resolutions. Both images are from the cross-validation of the general model where no local data is used during training. In the top image, there was no zooming and cropping data augmentation to mimic flying at different elevation heights. The result is that the model looks for birds at the exact size of the input training data, which are all smaller in size than this high resolution image. After training with augmented images, the model is able to adjust to moderate changes in input resolution. Ground truth annotations are shown in orange, predicted bird detections are shown in blue.

#### 9. Atlantic Seabirds from Seabird Watch

Seabird Watch is a project run by the University of Oxford and University College Cork. We have developed a Palearctic camera network, which has been collecting images of Black-legged Kittiwake (*Rissa tridactyla*) and guillemots (*Uria aalge* and *Uria lomvia*) since 2014. Our ground-based time-lapse cameras are located across the North Atlantic, and include sites in the UK, Ireland, Faroes, Iceland, Greenland and Svalbard. Images are annotated by volunteers as part of our citizen science project hosted on the Zooniverse platform (https://www.zooniverse.org/projects/penguintom79/seabirdwatch), as well as researchers who provide ‘gold standard’ annotations for comparison. For this study, only gold standard annotations have been used. We aim to investigate variables affecting seabird ecology and demography across large spatial and temporal scales. In particular, we aim to determine chick survival and breeding success and how this varies across species’ ranges; identify the causes of chick mortality; and record changes in the timing of breeding and how this is affected by environmental conditions. This dataset is the only non airborne image capture, but sufficiently similar due to its angle and distance.

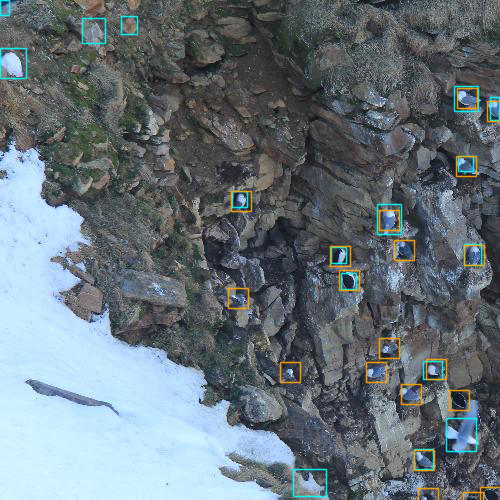
seabird watch

Figure S12. Sample test image from the Seabird Watch dataset of guillemots and kittiwakes from the North Atlantic. Predictions from the cross-validation model are in blue, ground truth annotations are in orange.

#### 10. Indian Ocean Seabirds

The Indian Ocean Seabird datasets were obtained on Pulu Keeling (North Keeling Island, 11.833 S. 96.82 E) and Christmas Island (10.45 S, 105.69 E) in the tropical central and eastern Indian Ocean.

Surveys on Pulu Keeling were conducted in November 2019 and February 2020 using a DJI Phantom 4 Pro V2 quadcopter at altitudes of between 50-55m. Although Pulu Keeling is a low lying coral atoll with relatively flat terrain much of the island is vegetated; Pemphis acidula shrublands which support Lesser Frigatebird colonies reach heights of 2-5 m, whereas Pisonia grandis forests which support Red-footed Booby colonies reach heights of 20-30 m. As a consequence flight height above vegetated canopy ranged from 20 to 50 m. Aerial imagery was captured using the standard in-built 20-megapixel camera for a Phantom 4 Pro with an 84° field-of-view. White balance, ISO, shutter speed and aperture were set to automatically adjust to variable light conditions, and focus was set to infinity. RPA flight speed was set to ~11 m/s, and image capture to a minimum of 75% front- and 70% side-overlap.

Surveys on Christmas Island were conducted in May 2021 using a DJI 4 RTK quadcopter. Christmas Island is a ‘high’ tropical island; the island is fringed by low sea cliffs behind which coastal terrace(s) extend inland to further cliffs (or steep slopes) that rise to the central plateau (max elevation 357 m). Given the complexity of the terrain, all surveys were flown in ‘terrain tracking mode’ at heights of between 50-60 m above ground. Aerial imagery was captured using the standard in-built 20-megapixel camera for a Phantom RTK with an 84° field-of-view. White balance, ISO, shutter speed and aperture were set to automatically adjust to variable light conditions with shutter speed priority set at > 1/1000 sec, and focus set to infinity. RPA flight speed ranged from 2.6 to 3.8 m/s, and image capture was set to a minimum of 85% front- and 80% side-overlap.

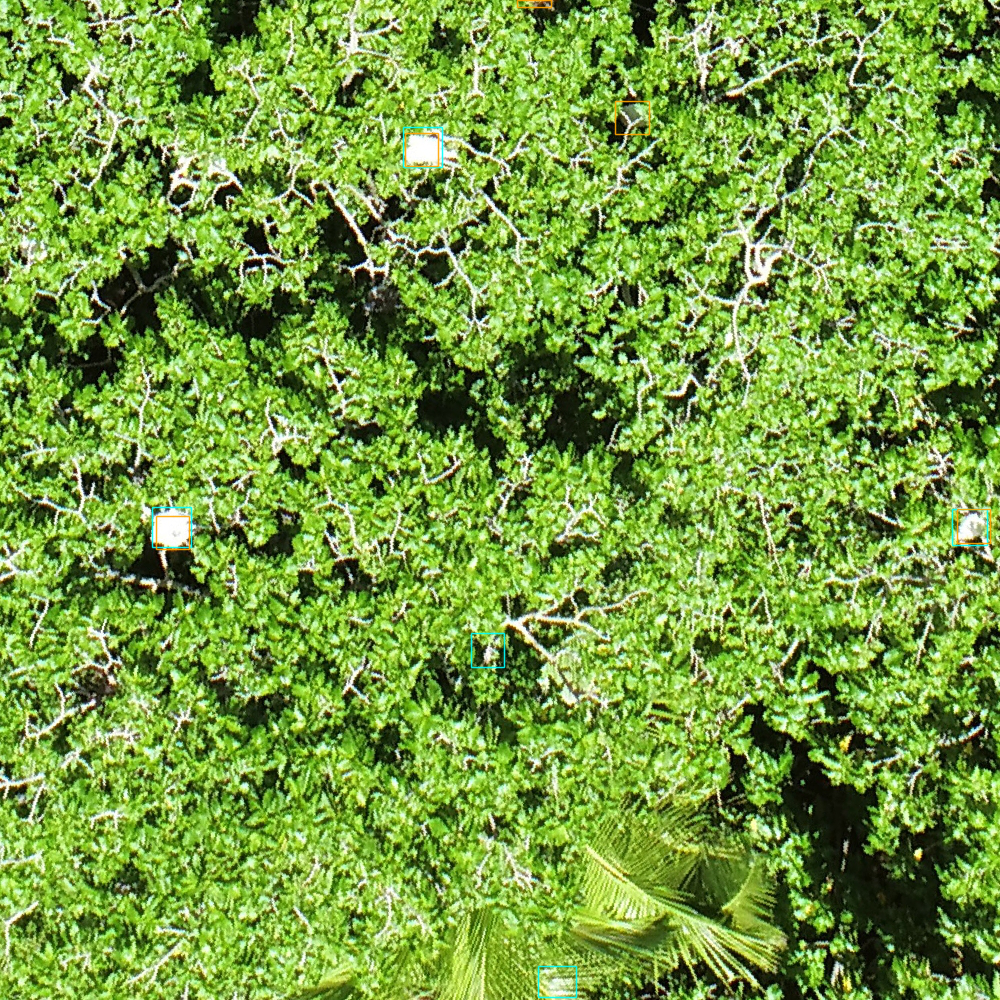

Figure S13. Sample test image from the seabird dataset from the Indian Ocean. Compared to the other forested datasets, the Indian Ocean dataset is overexposed and the birds are difficult to see. Predictions from the cross-validation model are in blue, ground truth annotations are in orange.

#### 11. Pelicans from Utah, USA

Yearly surveys of American White Pelicans (Pelecanus erythrorhynchos) from 2013 - 2020 were performed by the Utah Division of Wildlife Resources at Great Salt Lake, Utah using piloted aircraft and an observer taking photographs. The approximate altitude of flight is 800-1300ft and images were captured using a Canon 5D Mark II with a 35-350mm zoom lens. Surveys are conducted in late spring each year and all pelicans within the images are annotated by hand using ImageJ. There are additional bird species, such as gulls, which are not annotated within the images.

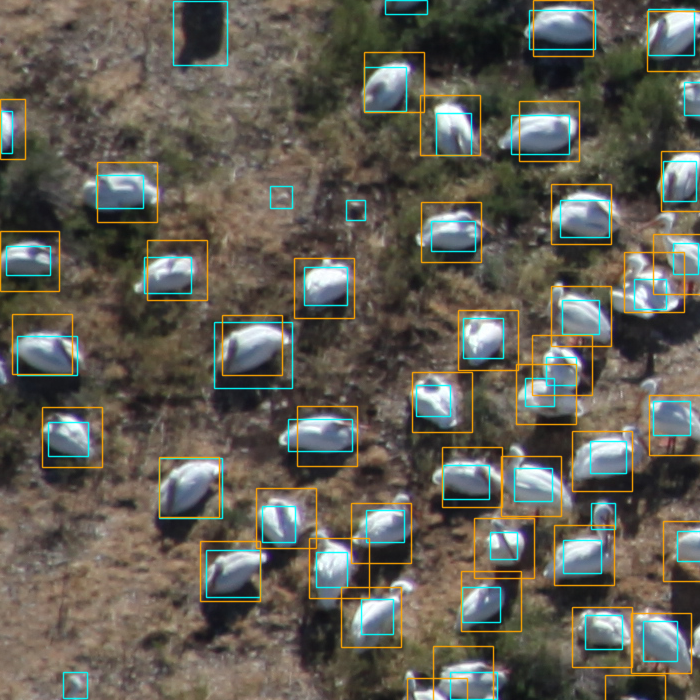

Figure S14. Sample test image from the pelican dataset from Utah, USA. Predictions from the cross-validation model are in blue, ground truth annotations are in orange.

#### 12. Cranes, Geese and Ducks from New Mexico, USA

Imagery of waterfowl was collected at two federal wildlife refuges in New Mexico as part of a collaborative project between the US Fish and Wildlife Service and the Center for the Advancement of Spatial Informatics Research and Education at the University of New Mexico.One survey was completed at Maxwell National Wildlife Refuge, located in northern New Mexico in a wetland environment adjacent to mixed shortgrass prairie, agricultural land, and pinon-juniper woodland at an average elevation of 1840m. Three flights were conducted December 15^th^, 2017 using a SenseFly EBEE+ drone equipped with a S.O.D.A. photogrammetric camera. The flights were conducted at an altitude of 60 m AGL, yielding a GSD of 1.4cm/px. Four surveys were completed at Bosque del Apache National Wildlife Refuge in seasonal wetland and riparian woodland adjacent to Chihuahuan desert and agricultural land at an average elevation of 1380m. Eleven flights were conducted across four field dates—November 6^th^, 7^th^, 13^th^, and 27^th^ 2018—using a DJI Mavic Pro 2 equipped with a Hasselblad L1D-20c sensor. Flight altitudes from 20 – 60 m AGL stepped in increments of 10m were tested on separate flights to determine the response of the waterfowl to the aircraft, yielding GSD 0.5cm/px - 1.4cm/px.A benchmark set of thirteen images was annotated by 18 biologists from the USFWS from August – September 2019 using the Labelbox proprietary platform. A twelve-species classification scheme was used, along with an “other” category for individual birds whose species was deemed indeterminate by the expert observers. Common species identified in the benchmark dataset include Sandhill cranes (*Antigone canadensis*), Canada geese (*Branta canadensis*), mallards (*Anas platyrhynchos*), and northern pintails (*Anas actua*).

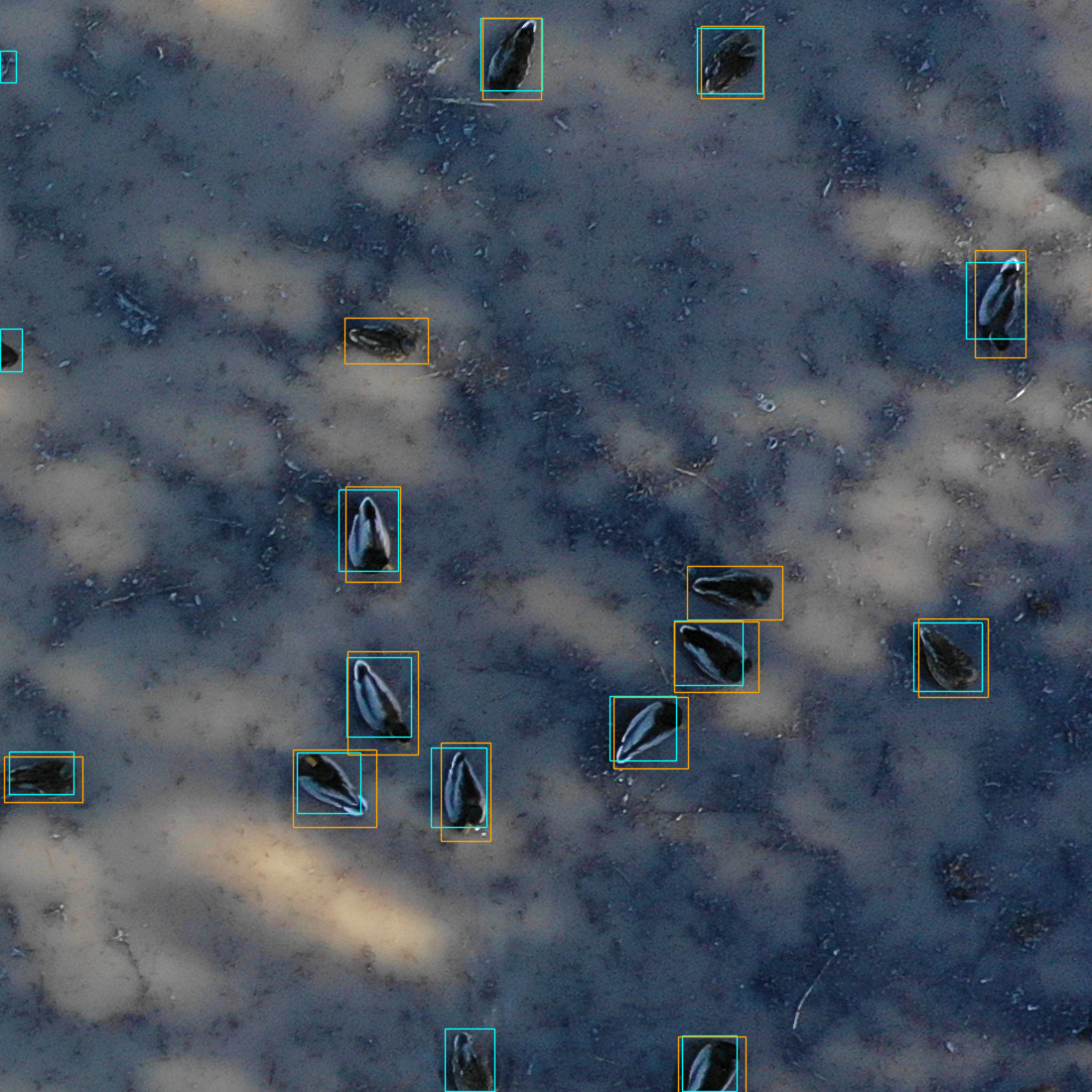

Figure S15. Example test image from cranes dataset from New Mexico. Predictions from the cross-validation model are shown in blue, ground truth annotations are in orange.

#### 13. Lake Michigan, USA

Imagery of waterfowl on Lake Michigan, USA was collected by the US Fish and Wildlife Service’s SEABirD sensor array during two sensor test flights on September 25, 2020. The waterfowl included mallards (*Anas platyrhynchos*), American white pelicans (*Pelecanus erythrorhynchos*) and gulls (species unidentified), among others. The SEABirD sensor array consists of seven mapping cameras integrated with a GPS/INS. Each camera has a CMOS Bayer color filter array (6464 x 4852 pixels) and 100mm lens with an F/4.0 aperture. The images were acquired from a Quest Kodiak aircraft at altitudes ranging from 600 to 1,000 ft AGL resulting in image ground resolutions of 0.6 to 1 cm/pixel. The test flights acquired 22,154 images, of which 634 were selected for ground truth annotation and used in this paper. A commercial data annotation firm generated 34,035 bounding boxes of birds from the selected images. The selected images exhibited backgrounds of open water, wetland, and shoreline with annotation counts ranging from one to over 1,000 birds per image.

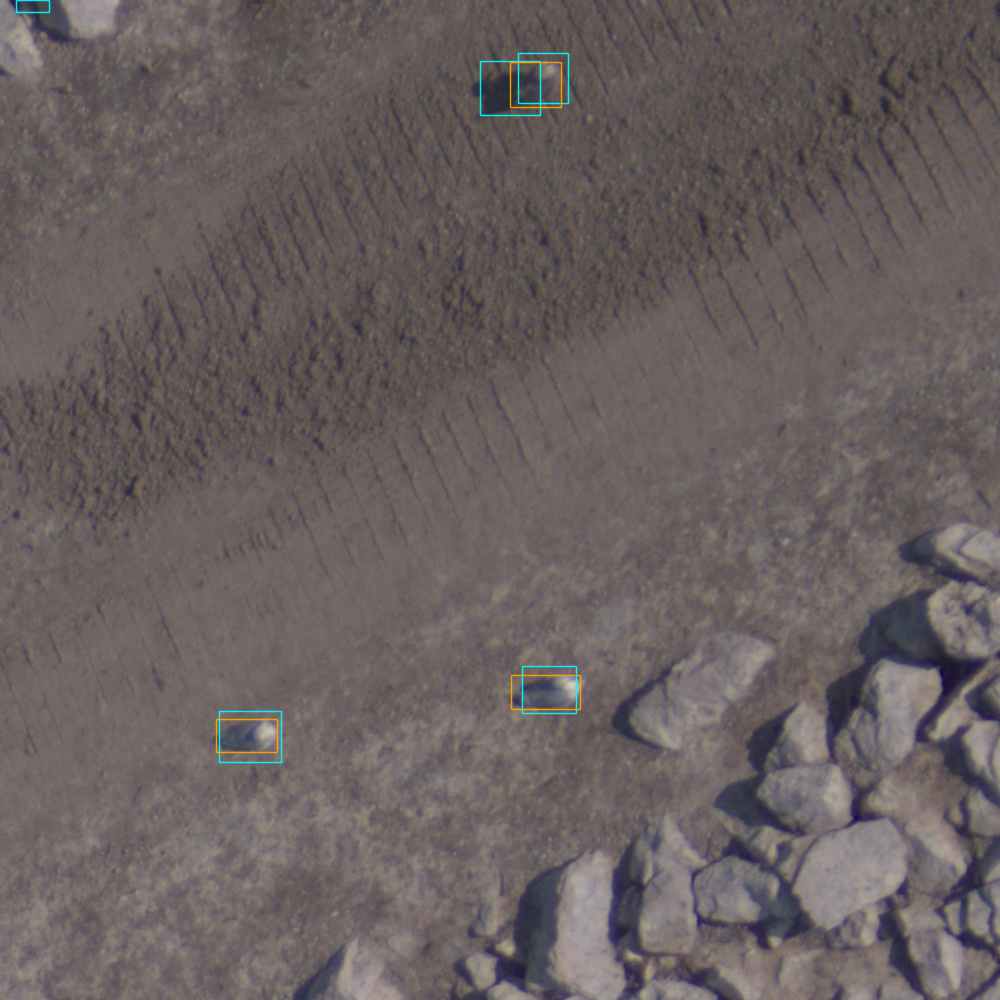

Figure S16. Sample test image from the Lake Michigan Dataset. Predictions from the cross-validation model are in blue, ground truth annotations in orange.

#### 14. Waterbirds from Poland

The study was carried out on colonial and gregarious species of waterbirds in northern Poland, in the years 2018 - 2021. Most of the observations were made in the lower course and estuary of a large lowland river – the Lower Oder Valley (site-centre location in decimal degrees: Longitude - 14.413200, Latitude - 53.085000). The observations were made in areas known for their importance for waterbirds, during the breeding season, migration, and the wintering period. A DJI Phantom drone (version 4 Pro V2.0) was used for the fieldwork. This is a remotely controlled quadcopter device with a total weight of 1,388 grams, equipped with a camera capable of taking both still photographs (max quality 5,472x3,648 pixels) and videos (max quality: 4,096 x 2,160 pixels), with the possibility of continuous tracking of unrecorded images transmitted to the display coupled with the remote control. Photos of the birds were taken from various heights in the range of about 20-80 meters. The photos used for this publication were taken of the following bird species: Black-headed Gull (*Chroicocephalus ridibundus*), Common Tern (*Sterna hirundo*), Gray Heron (*Ardea cinerea*), Mallard (*Anas platyrhynchos*), Mute Swan (*Cygnus olor*), Tufted Duck (*Aythya fuligula*), Greater Scaup (Aythya marila), Eurasian Wigeon (*Mareca penelope*), Eurasian Teal (*Anas crecca*), Northern Pintail (*Anas acuta*). The poland dataset covers a wide range of conditions and camera settings and does not represent a single project, but rather a collection of work over a period of years. Due to this, we did not include a test split, since there is no natural way to split images to create a coherent connection among images similar to the other datasets in this paper.

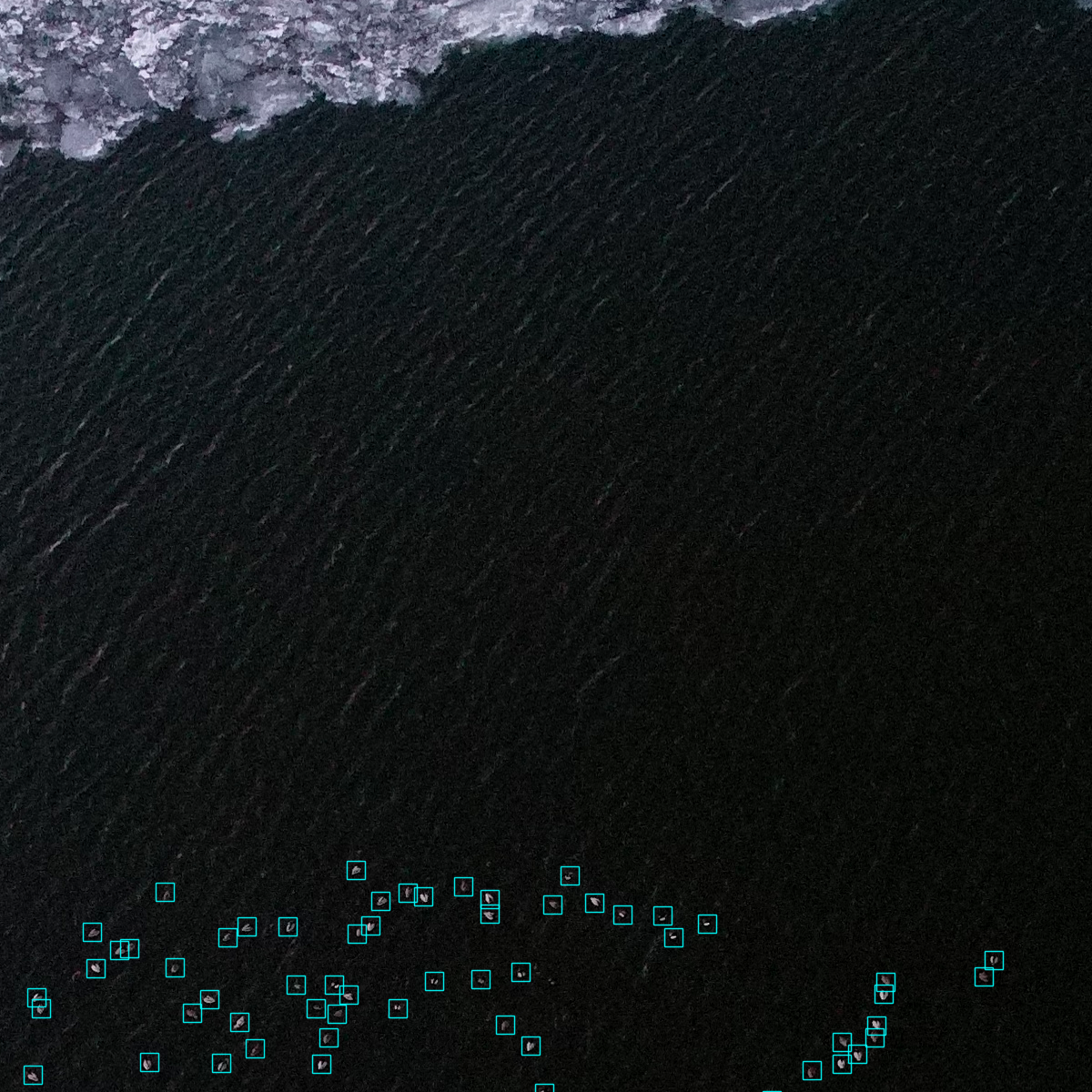

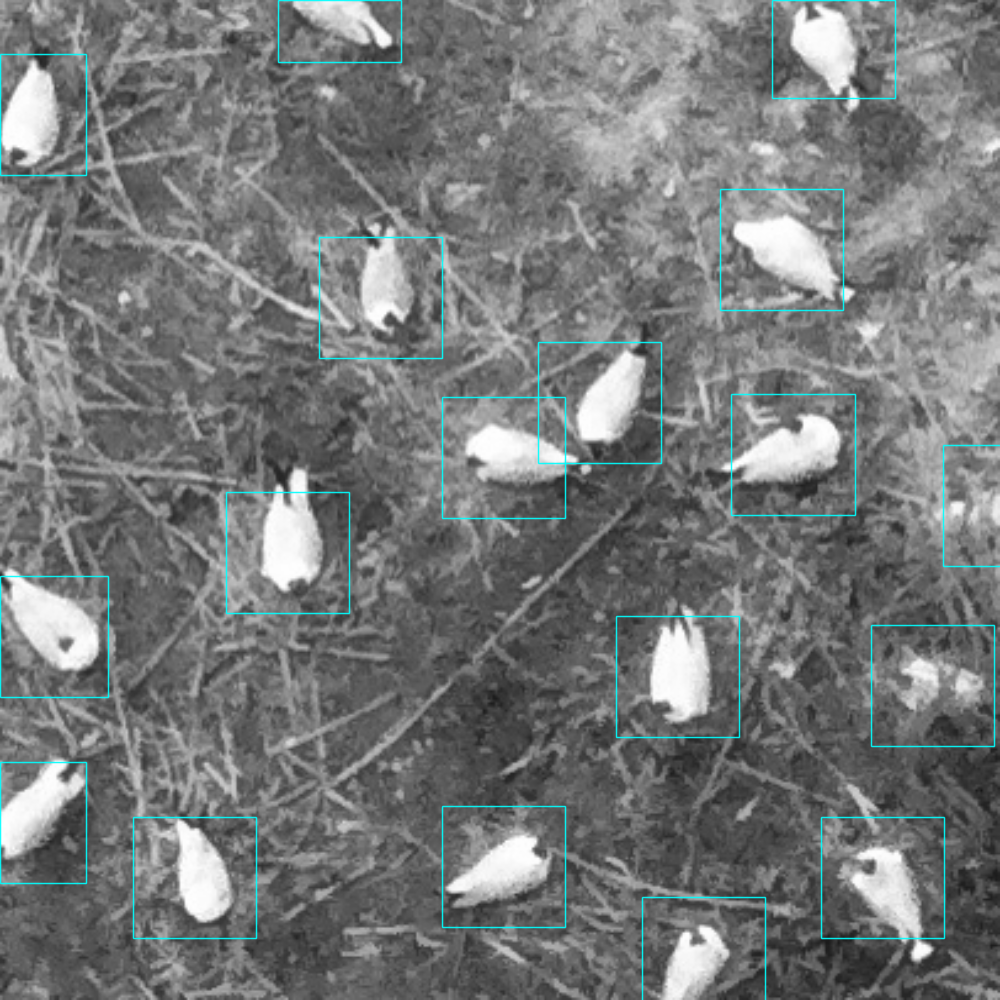

Figure S17. Sample training images from the Poland data. Because the dataset does not represent one project, it was not used in the cross-validation analysis.
